# Embryonic integrin signaling establishes a program of differential cell elongation in a growing proprioceptive organ

**DOI:** 10.64898/2026.11.07.765072

**Authors:** Praween Kumar Choubey, Adel Avetisyan, Lihi Amit, Rina T. Greenblatt Ben-El, Naomi Halachmi, Adi Salzberg

## Abstract

The ChO is a proprioceptive sensory organ that undergoes extensive morphological remodeling during larval growth. A key feature of this process is the dramatic elongation of cap cells, while adjacent ligament cells remain short. How these contrasting growth behaviors are established and coordinated is largely unknown. Using targeted RNAi screens of extracellular matrix (ECM) components, ECM receptors, and their intracellular effectors, we identified distinct molecular mechanisms controlling ChO architecture. We found that basement membrane components, including collagens and laminins, are required to maintain the characteristic straight morphology of cap cells, whereas Thrombospondin and Tiggrin contribute to the stability of cellular attachments.

Unexpectedly, we discovered that Thrombospondin and an αPS2βPS integrin signaling module, comprising Talin and the ILK–PINCH–Parvin complex, regulate the relative lengths of cap and ligament cells. Cell-type-specific knockdown demonstrated that this pathway acts within ligament cells to restrict their elongation. Disruption of integrin signaling causes ligament cells to elongate dramatically at the expense of cap cell extension, effectively reversing their normal growth behaviors. Importantly, this function is distinct from the role of integrin-associated proteins in maintaining cellular attachments. Temporal manipulation of integrin expression revealed that the short morphology of ligament cells is established during a critical developmental time window during embryogenesis, before the organ becomes functional. Restoring integrin expression after hatching fails to rescue the phenotype, indicating that embryonic integrin signaling establishes a persistent developmental state. Together, these findings reveal that ligament cell elongation is actively restricted by an embryonically established integrin-dependent mechanism, providing the structural basis for subsequent cap cell elongation and organ maturation.

## Introduction

Different cell types that constitute a multi-cellular organ often acquire markedly different morphologies and mechanical properties. Understanding how these differences arise and are maintained is essential for elucidating the mechanisms underlying the development of functional multicellular organs. The chordotonal organs (ChOs), the main proprioceptive organs of the *Drosophila* larva, provide a model for investigating this question (Caldwell et al., 2003). Each ChO comprises a sensory unit containing a ciliated neuron ensheathed by a glia-like scolopale cell, connected to the epidermis through a cap cell on the dendritic side and a ligament cell on the axonal side (Lai and Orgogozo, 2004)(Fig. 1). The best-characterized larval ChO is the lateral pentascolopidial organ (LCh5), a cluster of five ChOs in each abdominal hemisegment. The five cap cells are anchored to the epidermis through two cap attachment (CA) cells, while the five ligament cells share a single ligament attachment (LA) cell (Fig. 1) (Ghysen and Dambly-Chaudiere, 1989; Inbal et al., 2004). During larval locomotion, muscle contraction lead to cuticle deformations which alter the distance between the cap- and ligament-attachment sites and cause the organ to shorten (Hassan et al., 2019). Most of this shortening occurs in the cap cells, whereas the ligament cells hardly deform (Hassan et al., 2019). The strain generated by cap cell compression activates mechanosensitive channels at the dendritic tip, generating neuronal activity (Gong et al., 2004; Hassan et al., 2019; Kim et al., 2003).

**Figure 1.**
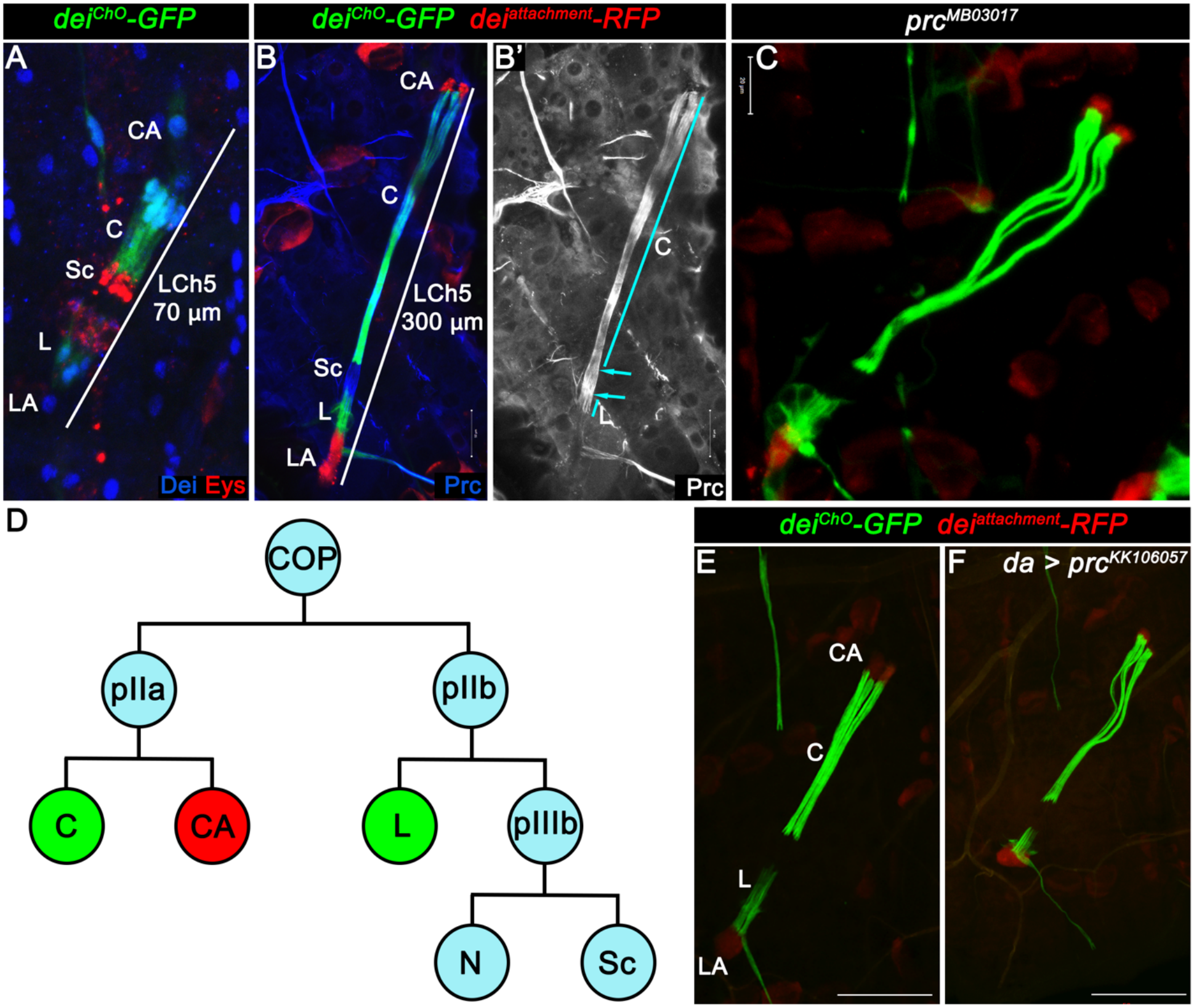
Differential responses of the LCh5 lineage-related cells to developmental stretching. (A) LCh5 organ of a stage 16 *dei^ChO^-GFP* embryo stained with anti-Eys (red) and anti-Dei (blue) to label the ligament (L), cap (C), scolopale (Sc) and attachment (LA, CA) cells. (B, B’) LCh5 organ of a 3^rd^ instar *dei^ChO^-GFP, dei^attachment^-RFP* larva stained with anti-Prc (blue). Prc localizes along the LCh5 organ, enwrapping the cap cells, the scolopale cells and part of the ligament cells above the neuronal cell bodies (B’). Arrows in B’ indicate the area occupied by Sc cells. Scale bar: 50 µm. (C) LCh5 organ of a 3^rd^ instar *dei^ChO^-GFP, dei^attachment^-RFP; prc^MB03017^* larva. Loss of Prc causes dramatic undulation of the cap cells. Scale bar: 20 µm. (D) Schematic representation of the asymmetric cell divisions that generate the five clonally related cells comprising a ChO. Cap and ligament cells express *dei^ChO^-GFP*, whereas cap attachment cells express *dei^attachment^-RFP*. (E) LCh5 organ of a 2^nd^ instar *dei^ChO^-GFP, dei^attachment^-RFP* larva. Scale bar: 50 µm. (F) Ubiquitous depletion of *prc* under the regulation of *da-Gal4* causes pronounced cap-cell undulation, which is already evident in 2^nd^ instar larvae. Scale bar: 50 µm.

The cellular organization of the LCh5 is established during embryogenesis, when asymmetric divisions of ChO precursor cells generate the neuron, scolopale, cap, ligament and CA cells (Fig. 1) (Brewster and Bodmer, 1995). Their subsequent differentiation is controlled by cell-type-specific transcriptional programs (Avetisyan et al., 2021; Campbell et al., 1994; Halter et al., 1995; Jones et al., 1995). Following cell specification, the ligament cells guide the ventral migration of the organ and recruit the LA cell once they reach their destination (Inbal et al., 2003; Inbal et al., 2004; Klein et al., 2010). LCh5 development extends beyond embryogenesis, as the organ undergoes extensive structural remodeling to accommodate larval growth. During this period, its total length increases from approximately 70 μm at the end of embryogenesis to more than 300 μm in 3^rd^ instar larvae (Halachmi et al., 2016). Remarkably, although both the cap and ligament cells are exposed to developmental stretching forces during organ elongation, their response is different. Cap cells elongate nearly 13-fold and ultimately account for 65–70% of the organ’s total length, whereas ligament cells undergo relatively little elongation (Halachmi et al., 2016). Previous work in our laboratory demonstrated that the transcription factors Dei and Sr, together with the ChO-specific α-tubulin αTub85E, are required to maintain normal cell-length proportions during larval growth (Hassan et al., 2018). Moreover, ligament cell-specific depletion of *dei* or *αTub85E* caused abnormal ligament elongation, suggesting that cell-intrinsic mechanisms may contribute to this asymmetric response to developmental stretching (Hassan et al., 2018). ChOs encounter substantial mechanical loads primarily after hatching; whether mechanisms underlying differential cell elongation are established during embryogenesis or emerge during larval development as an adaptation to mechanical stresses imposed on the organ as the larva grows, remains to be elucidated.

Beyond the regulation of differential cell elongation, maintaining the mechanical properties of cap cells and the stability of ChO attachments is essential for normal mech-anosensory function. The extracellular matrix (ECM) contributes to both requirements by providing mechanical support and mediating cell adhesion (Greenblatt Ben-El et al., 2017; Hassan et al., 2019). Our previous work demonstrated that Pericardin (Prc), a collagen IV-like protein associated with the cap-cell ECM, is required for normal ChO mechanosensation (Fig. 1). Loss of Prc reduced cap-cell elasticity and caused undulation. During muscle contraction, these cells underwent buckling instead of normal axial compression, resulting in impaired mechanosensory responses and uncoordinated larval locomotion (Hassan et al., 2019). Interestingly, Prc is produced by non-ChO cells, secreted into the extracellular environment, and subsequently incorporated into the ECM surrounding the organ (Chartier et al., 2002; Drechsler et al., 2013), highlighting the contribution of ECM components supplied by other tissues. The ECM also contributes to the stability of the LCh5 attachment sites. Thrombospondin (Tsp) accumulates at both cap–CA and ligament–LA junctions together with βPS integrin and its associated proteins and is required for stable cap–CA attachment (Greenblatt Ben-El et al., 2017; Halachmi et al., 2016). Moreover, depletion of βPS integrin within the ChO lineage disrupts cap–CA attachment and occasionally causes ligament-cell elongation (Greenblatt Ben-El et al., 2017; Hassan et al., 2018). These observations suggest that ECM-associated mechanisms may influence not only cell morphology and attachment stability but also the differential elongation of cap and ligament cells. However, the ECM components and intracellular mechanisms underlying these processes during larval growth remain poorly understood.

Here, we combined a targeted RNAi screen of ECM components, ECM receptors and their intracellular effectors with cell-type-specific and temporally controlled knockdown to investigate the mechanisms affecting LCh5 morphogenesis. Ubiquitous knockdown revealed that basement-membrane proteins and associated components are required for maintaining the straight morphology of cap cells, whereas Tsp and Tig contribute to cap–CA junction integrity. In addition, Tsp was required for ligament–LA attachment and for maintaining the relative lengths of cap and ligament cells. Among all genes examined, this latter function was shared only by components of an αPS2βPS– Talin–IPP signaling complex. Cell-type-specific manipulations revealed that this complex acts within ligament cells to restrict their elongation, thereby permitting cap cells to extend during larval growth. Temporal knockdown experiments further demonstrated that the requirement for integrin signaling in ligament cells is predominantly embryonic and that restoring its activity during larval development does not reverse the resulting defects. Together, these findings indicate that ligament cell length is actively constrained by an integrin-dependent mechanism established during embryogenesis, providing the developmental basis for the unequal elongation of cap and ligament cells during larval growth.

## Results

### Identification of ECM components required for LCh5 morphogenesis

Building on our previous discovery that the collagen IV-like ECM protein Prc is essential for cap cell mechanosensing and that its loss produces a striking LCh5 morphological phenotype (Fig. 1) (Hassan et al., 2019), we sought to identify additional ECM components required for LCh5 morphogenesis that may regulate the properties of cap cells or other cell types within the LCh5 organ. We therefore performed a targeted RNAi screen focusing on genes involved in ECM composition, assembly, and cell–ECM interactions. Candidate genes were selected through a combination of FlyMine (Lyne et al., 2007) searches using the terms *“ECM assembly”, “positive regulators of ECM assembly“* and *“ECM receptors”,* together with manual curation of genes from the published *Drosophila* matrisome (Davis et al., 2019) and additional transmembrane proteins likely to mediate ECM–cell or cell-cell interactions (Supplementary Table 1). The final screen comprised 135 genes, including 66 matrisome genes, representing 10.3% of the annotated *Drosophila* matrisome. For each of the 135 genes, we tested at least two independent, non-overlapping RNAi lines where available for a total of 320 RNAi lines screened. Every *Drosophila* collagen and laminin gene, which encode major structural components of basement membranes, was included in the screen (Supplementary Table 1).

To identify ECM proteins produced both locally and by distant tissues before their incorporation into the LCh5 ECM, RNAi was driven ubiquitously using *daughterless* (*da*)-Gal4. LCh5 morphology was assessed in 2^nd^ instar larvae using a dual fluorescent reporter system in which GFP labels cap and ligament cells (*dei^ChO^-GFP*) and RFP labels cap attachment, ligament attachment and tendon cells (*dei^attachment^-RFP*) (Halachmi et al., 2016). This strategy enabled rapid, high-throughput evaluation of LCh5 morphology in intact fixed larvae without the need for dissection or immunostaining (Fig. 1). The phenotypic consequences of each RNAi were scored in multiple larval segments and phenotype frequencies were compared with those observed in control larvae expressing *luciferase RNAi* under the same driver. An RNAi line was considered positive when the phenotype frequency exceeded that of the control and reached at least 15%, and a gene was considered a hit when at least two independent RNAi lines were positive.

To validate this screening approach, we examined *prc*. Ubiquitous *prc* knockdown using two independent RNAi lines induced pronounced cap cell undulations in second-instar larvae, affecting 52% (32/61) and 59% (26/44) of hemisegments, respectively, compared with only 5% (2/42) in controls (Fig. 1 and Table 1). Strikingly, LCh5-wide *prc* knockdown using *en*-Gal4 had little effect on cap cell morphology (3/64 organs). This is consistent with Prc being produced by non-LCh5 cells (Chartier et al., 2002) and highlights the importance of ubiquitous gene depletion for identifying ECM components that are essential for proper LCh5 morphogenesis.

### ECM components are required for the establishment of cap cell morphology

The first group of hits primarily affected cap-cell morphology, with phenotypes ranging from undulations to severe developmental abnormalities. Under normal conditions, cap cells exhibit a straight, elongated morphology and uniform *dei^ChO^-GFP* fluorescence along their entire length (Figs. 1 and 2A). This organization was largely preserved in control (*da* > *Luc RNAi*) larvae, with undulations observed in only 5% of LCh5 organs (2/42; Table 1).

**Figure 2.**
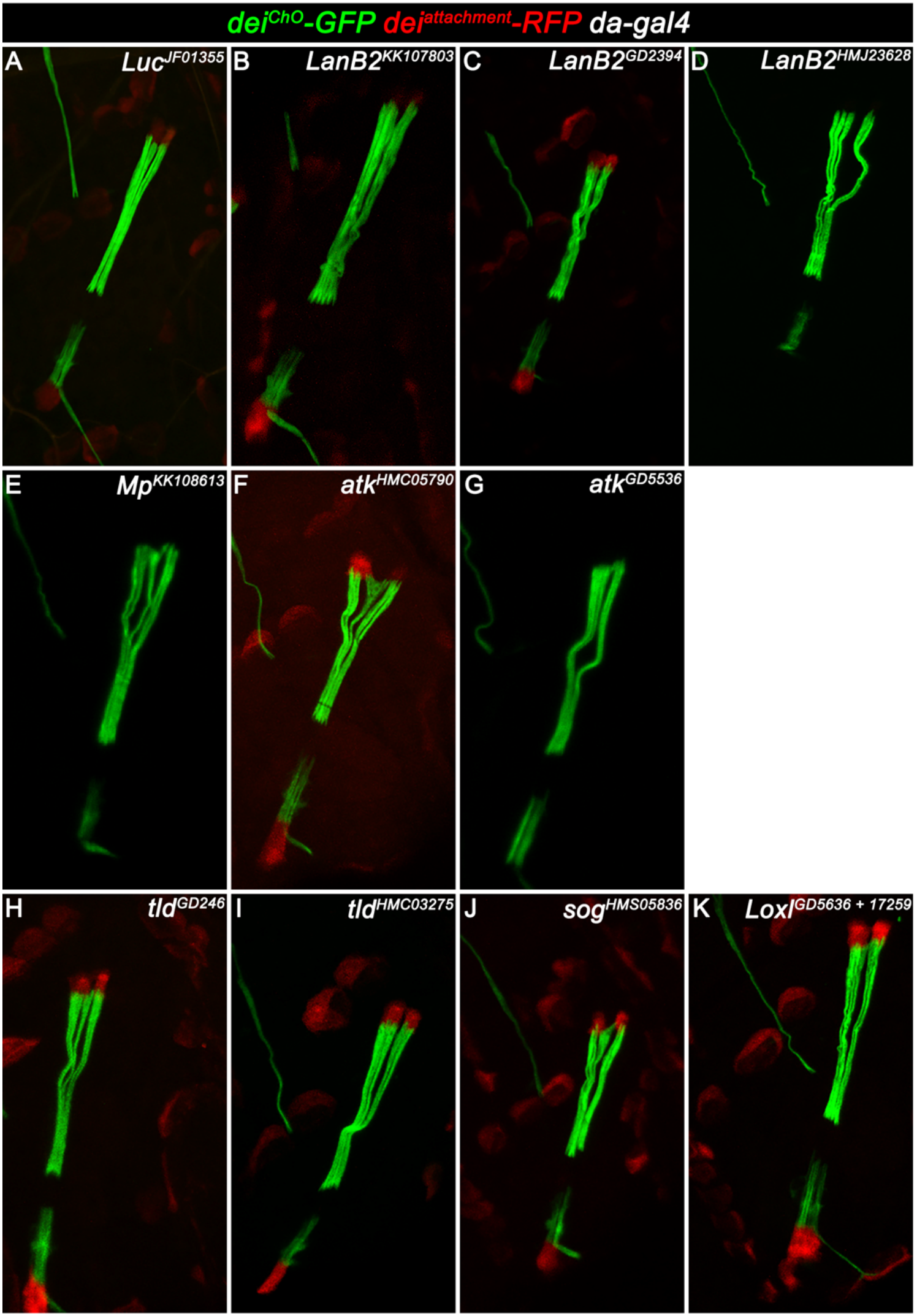
Basement membrane components are required to maintain the straight morphology of ChO cap cells. (A-K) LCh5 organ of a 2^nd^ instar *dei^attachment^-RFP; dei^ChO^-GFP, da-gal4* larva expressing either control RNAi (A) or RNAi targeting *LanB2* (B-D), *Mp* (E), *atk* (F, G), *tld* (H, I), *sog* (J) and *Loxl1* (K). Depletion of each of these ECM components causes cap-cell undulation.

Among basement membrane components, depletion of type IV collagen and collagen XV/XVIII produced distinct phenotypes. Knockdown of *viking* (*vkg*), encoding the α2 chain of type IV collagen, caused severe developmental defects, including late embryonic or early larval lethality. Surviving first-instar larvae displayed abnormal LCh5 positioning, shortened cap cells, and loss of *dei^ChO^-GFP* fluorescence. Depletion of *Col4a1* (*Cg25c*), encoding the α1 chain of type IV collagen, produced variable results, with one RNAi line causing pleiotropic defects and another showing no detectable phenotype (Table 1). In contrast, depletion of *Multiplexin* (*Mp*), the *Drosophila* homolog of collagen XV/XVIII, primarily affected cap-cell morphology, causing undulations in 23% and 33% of hemisegments with two independent RNAi lines (Fig. 2 and Table 1).

Laminin depletion also affected cap-cell morphology. Three *LanB2* RNAi lines caused undulations in 23–42% of LCh5 organs, whereas two additional lines resulted in embryonic lethality. Similarly, *LanA* depletion induced cap-cell undulations. Two *LanB1* RNAi lines caused embryonic lethality, while a third produced no detectable phenotype (Table 1). These findings implicate several basement membrane components in maintaining normal cap-cell morphology.

The screen also identified proteins involved in extracellular matrix processing. Depletion of *tolloid* (*tld*), encoding an extracellular metalloprotease (Marqués et al., 1997), caused cap-cell undulations in 24–39% of LCh5 organs with three independent RNAi lines. Knockdown of its paralog, *tolkin* (*tok*) (Finelli et al., 1995), produced a similar phenotype, with frequencies ranging from 19% to 28%. Depletion of *short gastrulation* (*sog*), encoding a secreted substrate of Tld and Tok, caused undulations in up to 27% of organs, while knockdown of *Loxl1*, encoding a lysyl oxidase implicated in ECM cross-linking (Kim et al., 2014; Molnar et al., 2003), produced a comparable phenotype (17–26%; Fig. 2 and Table 1).

Unexpectedly, depletion of *artichoke* (*atk*), encoding a leucine-rich repeat ECM protein secreted by cap cells into the dendritic cap (Andrés et al., 2014), also caused pronounced cap-cell undulations, affecting 43% (9/21) and 25% (6/24) of hemisegments with two independent RNAi lines (Fig. 2 and Table 1). Thus, proteins with distinct extra-cellular localizations and functions contribute to maintaining the characteristic straight morphology of cap cells.

### Junction-associated ECM components regulate cell attachment and cell elongation

A second class of ECM components was associated with defects localized to the cap–CA junction, including cap cell thinning near the CA interface and detachment of cap cells from the CA cell (Fig. 3). Neither defect was observed in control larvae (*da > Luc RNAi*; Table 1). Depletion of *tiggrin* (*tig*) induced cap cell thinning near the CA interface with two of four independent RNAi lines, affecting 44% (11/25) and 26% (7/27) of LCh5 organs, respectively (Fig. 3 and Table 1). Although *tig* knockdown also produced occasional cap–CA detachments (3/29, 2/27, and 1/25 hemisegments with three RNAi lines), their frequency did not differ significantly from controls (Table 1). In contrast, depletion of *thrombospondin* (*tsp*) caused cap–CA detachment with both RNAi lines, affecting 17% (7/42) and 19% (13/70) of organs, respectively (Fig. 3 and Table 1). Notably, cap cells that remained attached generally retained their normal morphology (Fig. 3). In addition to its effects on the cap–CA junction, knockdown of *tsp* caused ligament cell detachment from the LA cell with both RNAi lines, affecting 17% (7/42) and 21% (15/70) of hemisegments, respectively, a phenotype not observed in controls (Fig. 3 and Table 1). Both Tig and Tsp are established components of *Drosophila* myotendinous junctions (Bunch et al., 1998; Chanana et al., 2007; Fogerty et al., 1994; Greenblatt BenEl et al., 2017; Subramanian et al., 2007; Vanhoutte et al., 2016), and our data raise the possibility that the cap–CA and ligament-LA interface contains a similar specialized adhesive ECM, involving Tsp and potentially Tig, that contributes to junctional stability.

**Figure 3.**
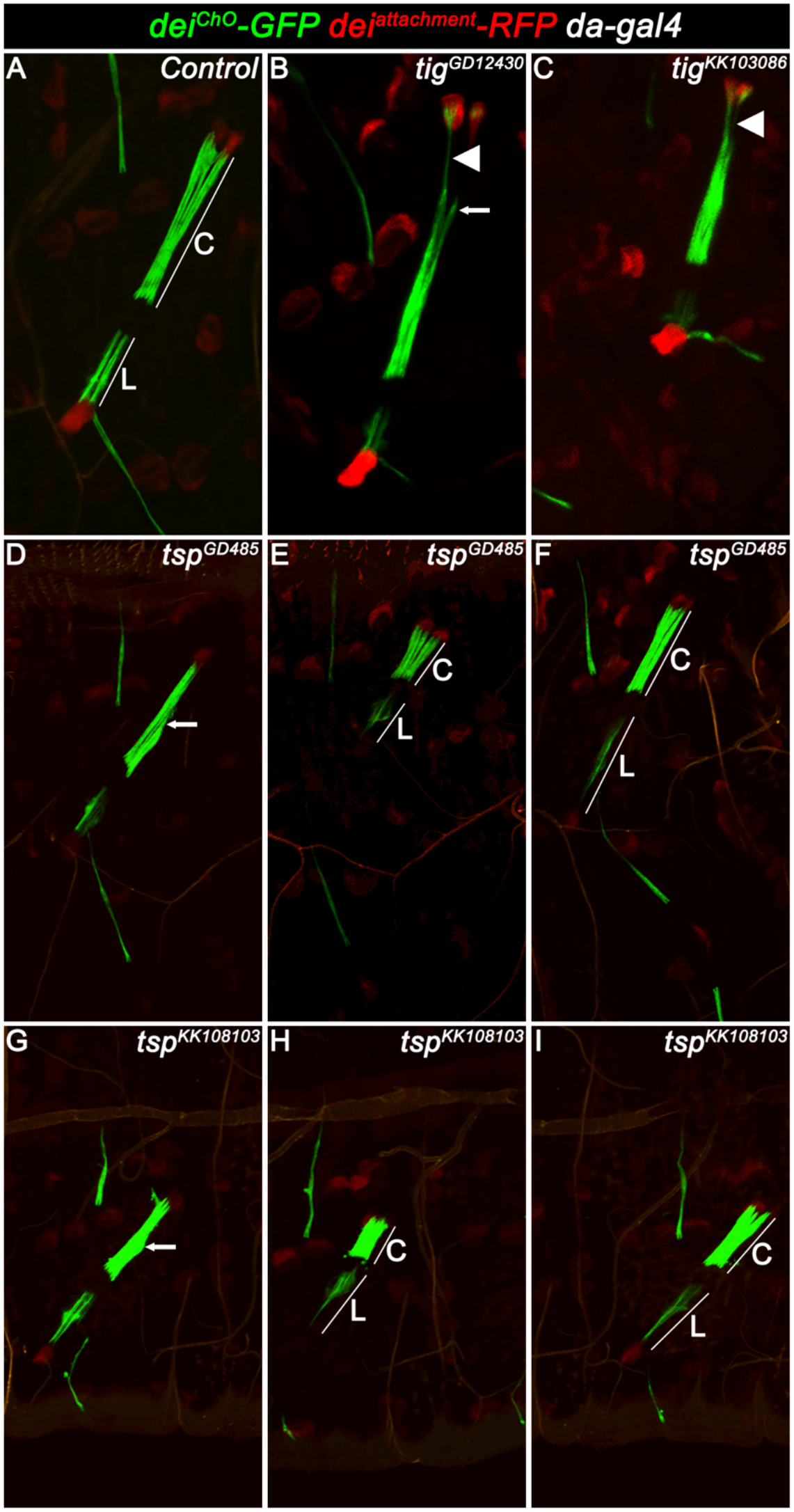
Tsp affects not only stability of the LCh5 junctional ECM, but also regulates ligament-cell elongation. (A-I) LCh5 organ of a 2^nd^ instar *dei^attachment^-RFP; dei^ChO^-GFP, da-gal4* larva expressing either control RNAi (A) or RNAi targeting *tig* (B, C) and *tsp* (D-I). (B, C) Ubiquitous depletion of *tig* resulted in pronounced thinning of cap cells near the cap-CA interface (arrowheads). Cap-cell detachment from the CA was rarely observed (arrow). (D-I) Ubiquitous depletion of *tsp* caused frequent cap-cell detachment (arrows in D and G), ligament-cell detachment from the LA cell (E, H) and abnormal elongation of ligament cells accompanied by decreased cap-cell length (F, I).

Strikingly, *tsp* depletion also revealed an unexpected role for the ECM in regulating the relative lengths of LCh5 cells. *tsp* depletion caused pronounced ligament-cell elongation accompanied by reduced cap-cell length (Fig. 3). To further characterize this phenotype, we measured the lengths of the cap cell and ligament-plus-sensory-unit (L+SenU) compartment in each hemisegment of control and *da > tsp RNAi* larvae and calculated the ratio of cap cell length to L+SenU length (C/[L+SenU]). Because loss of ligament anchorage could secondarily affect cell lengths, we excluded LCh5 organs exhibiting ligament–LA detachment from subsequent analyses. Knockdown with one RNAi line significantly reduced the C/(L+SenU) ratio, with 57% (20/35) of hemisegments exhibiting a length ratio falling below the threshold established from control organs, compared with none of the 42 control organs (Fig. 3 and Table 1 and 2). Relative to size-matched controls, the L+SenU compartment was 19% longer, whereas cap cells were 13% shorter. Moreover, pronounced ligament elongation was observed in 46% (16/35) of LCh5 organs. A second RNAi line produced a weaker but statistically significant shift in elongation ratios, with a reduced cap/L+SenU length ratio in 14% (7/49) of hemisegments and ligament elongation in 10% (5/49) (Fig. 3 and Table 1). The effect on LCh5 cell elongation pattern was remarkably specific to *tsp* depletion: among 132 other RNAi lines and 5,854 LCh5 organs examined in this part of the screen, a reduced cap/(L+SenU) ratio was observed in only five organs, from four RNAi lines targeting unrelated genes (Table 1). Thus, in addition to its role in stabilizing cell–cell attachments, Tsp emerges as a candidate regulator of the differential elongation of cap and ligament cells.

The distinct phenotypes identified in the screen prompted us to examine the distribution of the corresponding ECM components. Endogenously GFP-tagged Vkg, LanA and LanB1 surrounded the entire LCh5 organ, with continuous localization along the surfaces of its cells (Fig. 4). In contrast, Prc distribution was restricted to the region covering the cap cells, the scolopale cells and the ligament cells enwrapping the dendrites (Fig. 1). Tsp, on the other hand, was highly enriched at the cap–CA, ligament–LA, and ligament– scolopale junctions and was also detected as puncta along the cap-cell surfaces (Fig. 4) as was previously described (Greenblatt Ben-El, 2017). These localization patterns were consistent with the distinct phenotypic effects of their depletion: perturbation of collagen IV and laminin components primarily affected cap-cell morphology, whereas Tsp depletion compromised junctional integrity and altered the relative lengths of cap and ligament cells.

**Figure 4.**
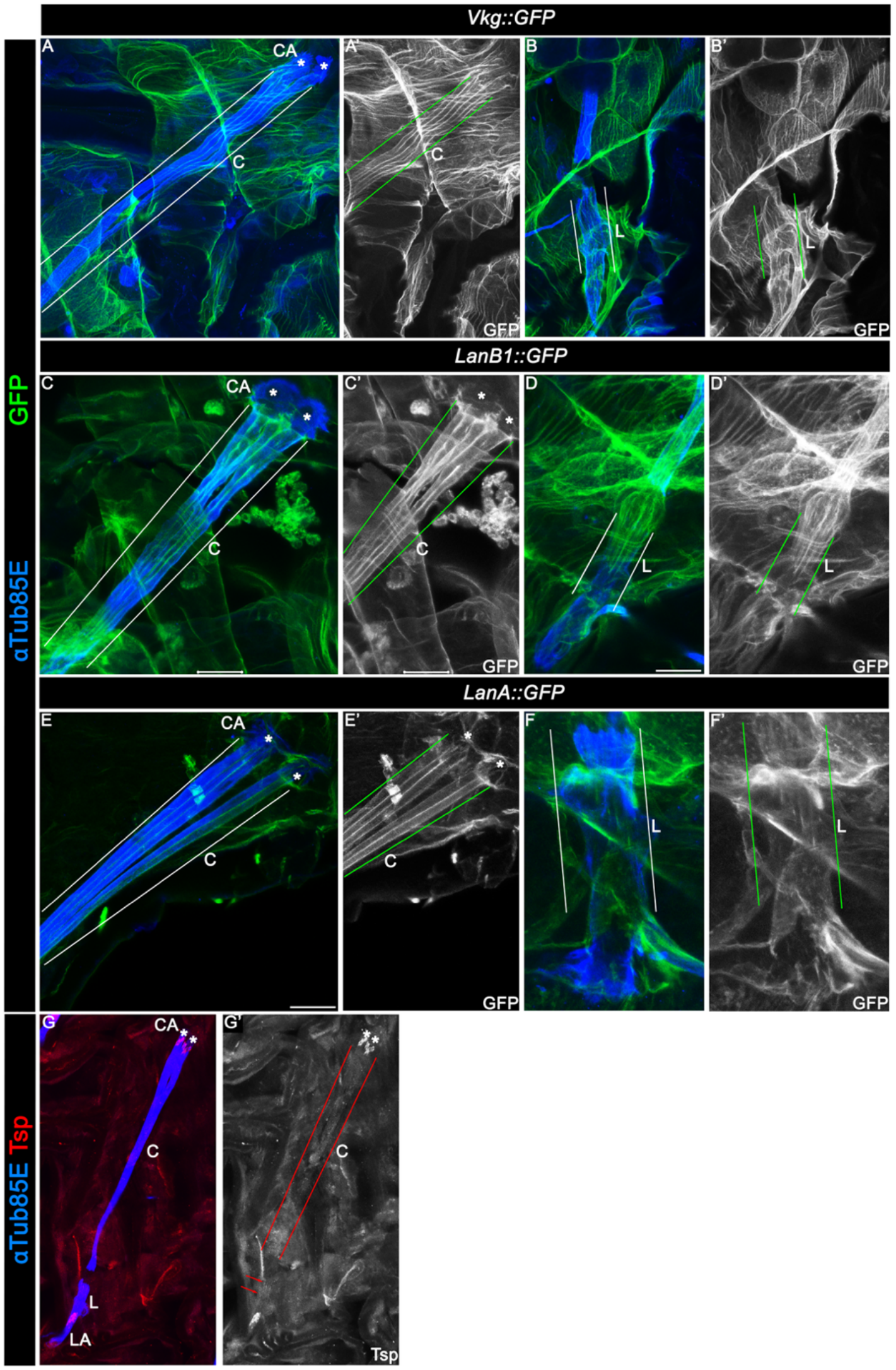
Localization of Vkg, LanB1, LanA and Tsp in the LCh5 organ. (A-F) LCh5 organ of 3^rd^ instar *Vkg::GFP* (A-B), *LanB1::GFP* (C-D) and *LanA::GFP* (E-F) larvae stained with anti-aTub85E (blue). All three proteins localize along the entire LCh5 organ. Scale bar: 20 µm. (G, G’) LCh5 organ of a wildtype 3^rd^ instar larva stained with anti-aTub85E (blue) and anti-Tsp (red). Tsp is enriched at the cap-CA, ligament-LA and ligament-Sc (red arrows in G’) interfaces and is also distributed in a punctate pattern along the cap cells.

Together, these findings indicate that LCh5 morphogenesis depends on functionally distinct extracellular environments: a basement membrane containing collagens and laminins that supports cap cell morphology, and a specialized junction-associated ECM that maintains stable connections between LCh5 accessory and attachment cells. Most unexpectedly, *tsp* depletion revealed a marked alteration in cap and ligament cell elongation, suggesting that it plays a role in maintaining the correct proportions of cap versus ligament cell lengths.

### Integrin signaling within LCh5 cells maintains the cap–CA junction and restricts ligament cell elongation

Because ECM components exert many of their effects through cell-surface receptors, we next asked whether such receptors are required within LCh5 cells for organ morphogenesis. Candidates were selected as described above and included established ECM receptors together with other transmembrane proteins, including members of the cadherin superfamily. Intracellular effectors of receptors identified as hits in the initial screen were subsequently examined in a second round (Supplementary Table 1). In total, 170 RNAi lines targeting 90 genes were screened. Because these proteins were expected to act within LCh5 cells, RNAi was driven throughout the LCh5 lineage using *engrailed* (*en*)-Gal4.

Reduced Integrin expression produced two distinct classes of morphological defects. The first affected cap cell morphology and attachment to the CA cell, consistent with our previous observation that integrin complexes are enriched at the cap–CA interface (Greenblatt Ben-El et al., 2017; Hassan et al., 2018). Affected cap cells exhibited pronounced thinning, particularly near their attachment to the CA cell, and, in more severe cases, detached from the CA cell altogether (Fig. 5). In control larvae (*en* > *Luc RNAi*), cap cell thinning was never observed (0/47 hemisegments), whereas cap–CA detachment occurred at low frequency (2/47 organs, 4%).

**Figure 5.**
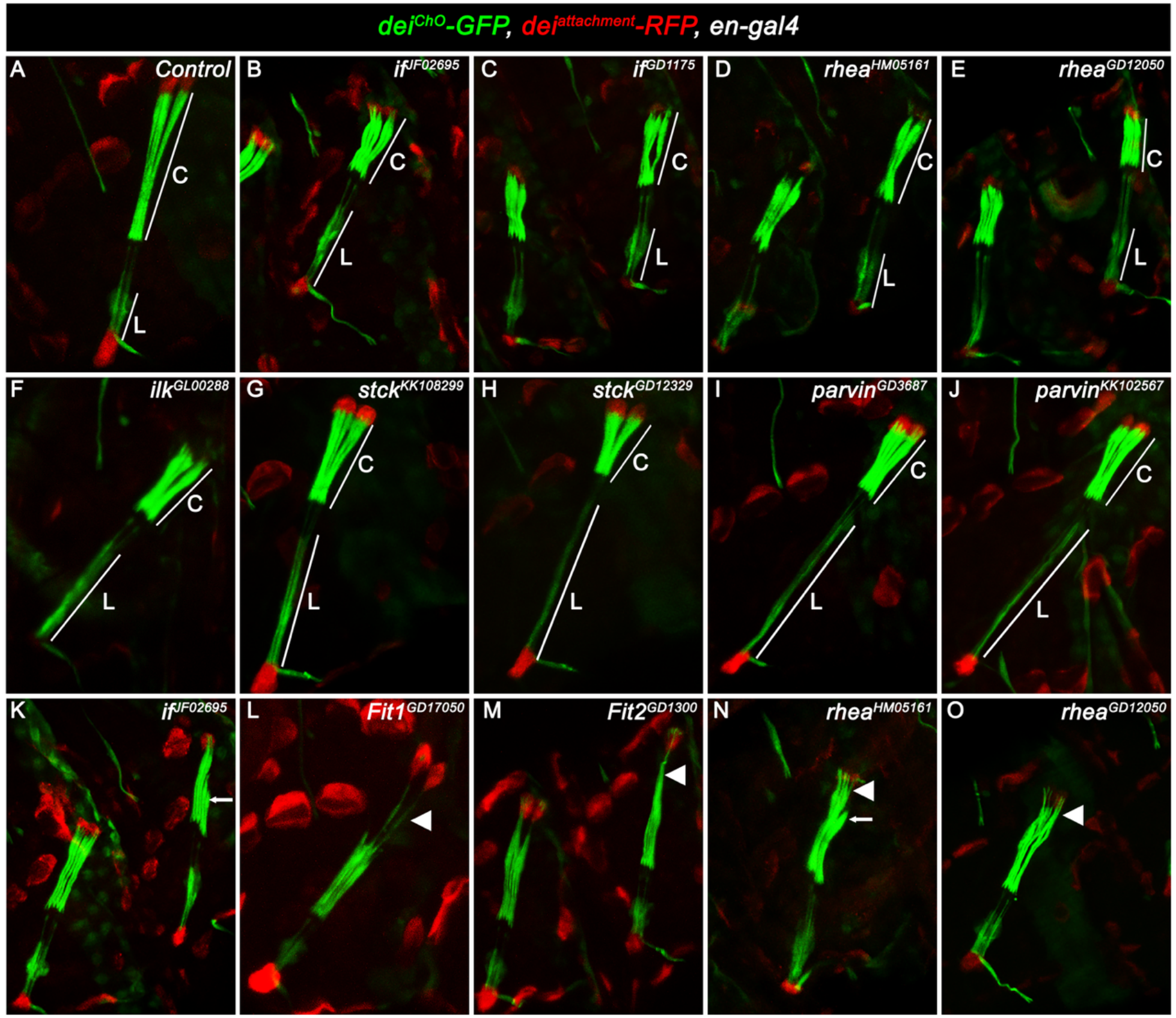
Integrin signaling is required for the correct pattern of differential cell elongation within the LCh5 organ and cap-CA junction stability. (A) LCh5 organ of a 2^nd^ instar *dei^ChO^-GFP, dei^attachment^-RFP, en-gal4* larva expressing control RNAi. (B-J) LCh5-wide depletion of *if* (B, C), *rhea* (D, E), *ilk* (F), *stck* (G, H) and *parvin* (I, J) resulted in abnormal elongation of ligament cells at the expense of cap cells. (K-O) LCh5-wide depletion of *if* (K) and *rhea* (N, O) also caused thinning of cap cells near the cap–CA junction (arrowheads) and cap-cell detachment (arrows). Cap-cell thinning was also observed following LCh5-wide depletion of *Fit 1* and *Fit 2* (L and M, respectively).

Knockdown of *inflated* (*if*), encoding the αPS2 integrin subunit, had only a limited effect on cap–CA junction integrity. Of the six RNAi lines tested, one induced cap cell thinning in 16% (3/19) of LCh5 organs, whereas cap–CA detachment observed with three other lines (5–6%, one organ per line) did not differ from controls (Fig. 5 and Table 3). In contrast, two *if* RNAi lines caused cap cell undulations, affecting 48% (13/27) and 24% (6/25) of LCh5 organs, respectively (3/47 in controls). Knockdown of *myospheroid* (*mys*), encoding the βPS integrin subunit, produced similar morphological defects (Fig. 5 and Table 3). By contrast, depletion of *multiple edematous wings* (*mew*), encoding the αPS1 integrin subunit, produced no significant morphological abnormalities with either of two independent RNAi lines (Table 3). Although the junctional defects following αPS2 depletion were mild and observed with only one RNAi line, the phenotypes associated with integrin perturbation, together with those observed following *tsp* and *tig*, are consistent with a possible role for integrin-mediated cell–ECM interactions at the cap–CA interface.

The second integrin-associated phenotype was of particular interest because, among the ECM genes examined in our screen, only *tsp* depletion had produced a similar alteration in the relative lengths of cap and ligament cells. In the integrin receptor screen, this phenotype was restricted to depletion of βPS and αPS2 integrins. We quantified these changes using the same approach as for *tsp* depletion, assessing the C/(L+SenU) ratio and L+SenU length relative to size-matched controls (Fig. 5 and Table 4). Reduced ratios and ligament elongation were defined as values falling below the 2.5th or above the 97.5th percentile of the corresponding size-specific control distributions, respectively.

Knockdown of *if* (αPS2) with two independent RNAi lines markedly reduced the mean C/(L+SenU) ratio with 93% (13/14) and 100% (20/20) of LCh5 organs falling below the threshold (Fig. 5 and Table 4). These effects remained evident after accounting for organ size, with 10/14 and 13/20 hemisegments falling below the size-specific threshold, compared with only 1/45 in controls. Relative to size-matched controls, the L+SenU compartment was 11% and 14% longer, whereas cap cells were 12% and 13% shorter, respectively. However, significant ligament cell elongation was detected only with the second line (6/20 hemisegments, compared with 1/14 for the first line). Three additional *if* RNAi lines produced milder reductions in the C/(L+SenU) ratio affecting 24–42% of the organs. However, none showed a significant effect relative to size-matched controls (1/17, 1/16, and 0/12 hemisegments below the size-specific threshold). The remaining line produced no detectable phenotype (Table 4).

Depletion of *myospheroid* (*mys*), βPS integrin subunit, produced a similar alteration in cell-length proportions (Fig. 5 and Table 3). In contrast, knockdown of *mew*, αPS1 integrin subunit, had no detectable effect with either of two independent RNAi lines (Table 4). This α-subunit specificity parallels the established requirement for αPS2, rather than αPS1, in Tsp-dependent myotendinous junction formation in *Drosophila* (Chanana et al., 2007). Together with the similar alterations observed following *tsp* depletion, these findings suggest that Tsp and αPS2βPS integrins may contribute to regulating the relative dimensions of cap and ligament cells.

We next investigated which intracellular components of integrin-associated complexes contribute to the two morphological phenotypes identified in the receptor screen. Depletion of *rhea*, encoding Talin (Lu et al., 2022), affected both cap–CA junction integrity and the relative lengths of cap and ligament cells. Cap cell thinning was observed with all four independent RNAi lines, affecting 9/14, 11/14, 7/21, and 3/17 LCh5 organs, respectively (Fig. 5 and Table 2). One line also caused pronounced cap–CA detachment in 44% (7/16) of hemisegments, whereas detachment frequencies with the remaining three lines did not exceed control levels. Three *rhea* RNAi lines also reduced the mean C/(L+SenU) ratio (Fig. 5 and Table 3). Because *rhea*-depleted organs were substantially shorter than controls, we also evaluated the lengths of the cap cell and the L+SenU region relative to total organ length. The reduction in the C/(L+SenU) ratio remained significant for two lines, with 16/21 and 6/17 organs falling below the size-specific control threshold, whereas the third line showed no significant effect (1/14). However, ligament elongation was rare, occurring in no more than two hemisegments per line. Thus, Talin depletion affects both cap–CA junction integrity and the relative dimensions of the cap and ligament cells.

In contrast, depletion of components of the integrin-linked kinase (ILK)–PINCH– Parvin (IPP) (Górska and Mazur, 2022) complex produced a particularly pronounced alteration in LCh5 cell-length proportions, with relatively little effect on cap–CA attachment. Knockdown of *stck*, encoding PINCH, reduced the mean C/(L+SenU) ratio with two independent RNAi lines, with all analyzed hemisegments falling below the threshold (Fig. 5 and Table 4). Unlike *if* and *rhea* depletion, *stck* knockdown caused a dramatic increase in L+SenU length, which was 1.8-fold greater than in size-matched controls for both lines, with every LCh5 organ exceeding the size-specific control range. A third RNAi line produced a milder phenotype, affecting 14/22 hemisegments and causing a 1.4-fold increase in L+SenU length (Table 4). Similarly, depletion of *parvin* with two independent RNAi lines reduced the mean C/(L+SenU) ratio in all analyzed LCh5 organs. These changes were accompanied by 1.8- and 1.6-fold increases in L+SenU length, respectively, relative to size-matched controls (Fig. 5 and Table 4). Knockdown of *Ilk*, the third component of the IPP complex, produced a similar elongated-ligament/shortened-cap phenotype with one of two RNAi lines tested (Fig. 5 and Table 3). Despite these pronounced changes in compartment proportions, depletion of *stck* or *parvin* rarely affected cap–CA junction integrity, with cap cell thinning or detachment observed in no more than two hemisegments per RNAi line (Table 4).

Depletion of *Vinculin* (*Vinc*), encoding a Talin-binding adaptor protein (Atherton et al., 2016; Li et al., 2025), produced a considerably milder phenotype. Two of five RNAi lines significantly reduced the size-adjusted C/(L+SenU) ratio, with 5/22 and 7/17 hemisegments falling below the size-specific threshold. These lines also caused ligament elongation in 4/22 and 6/17 hemisegments, whereas the remaining three lines had no detectable effect (Table 4).

Together, these findings reveal distinct phenotypic consequences of perturbing intracellular integrin-associated components. Whereas Talin depletion affects both cap– CA junction integrity and compartment proportions, disruption of the IPP complex preferentially produces pronounced elongation of the L+SenU compartment. These results highlight a particularly important contribution of IPP complex components to maintaining the normal relative dimensions of LCh5 cells.

Depletion of Kindlin-family integrin co-activators produced a distinct phenotypic profile (Böttcher et al., 2009; Lu et al., 2022), predominantly affecting cap cell morphology and attachment rather than the relative lengths of cap and ligament cells. Knockdown of *Fermitin 1* (*Fit1*) caused pronounced cap cell thinning in 16/17 hemisegments, accompanied by cap–CA detachment in 5 (29%). Similarly, depletion of *Fermitin 2* (*Fit2*) induced cap cell thinning in all 21 LCh5 organs examined with one RNAi line and in 12/26 with a second line (Fig. 5 and Table 3). Both *Fit2* RNAi lines also produced cap cell undulations, affecting 41% (9/22) and 23% (6/26) of organs, respectively. In contrast to the pronounced changes in compartment proportions following depletion of IPP-complex components, *Fit1* and *Fit2* knockdown did not reduce the C/(L+SenU) ratio (Fig. 5 and Table 4). Together, these findings suggest that Kindlin-family proteins contribute primarily to maintaining cap cell morphology and junctional integrity, with no effect on the relative lengths of the cap and ligament cells.

The distinct phenotypes resulting from depletion of different integrin-associated proteins raised the possibility that junctional integrity and the regulation of cap-to-ligament proportions represent separable aspects of LCh5 morphogenesis. To investigate this, we examined the co-occurrence of these defects in individual hemisegments across six integrin-pathway genotypes (*if*, *rhea*, *stck*, *parvin*, *Fit1*, and *Fit2*). Among 304 hemisegments analyzed, 73 exhibited cap cell thinning, of which only 15 (21%) also had a reduced C/(L+SenU) ratio. In contrast, a reduced ratio was observed in 69% (159/231) of LCh5 organs without cap cell thinning. A similar inverse association was observed for cap–CA detachment: only 33% (7/21) of hemisegments exhibiting detachment had a reduced C/(L+SenU) ratio, compared with 59% of those without detachment. Thus, junctional defects were significantly less likely to coincide with altered cell-length proportions, consistent with the differential involvement of integrin-associated proteins in these two aspects of LCh5 morphogenesis.

Because integrin-mediated adhesion is closely linked to the actin cytoskeleton (Lad et al., 2007), we extended our screen to genes involved in cytoskeletal organization and cell polarity to identify potential downstream effectors (Supplementary Table 1). None of the tested genes reproduced the cap–CA junctional defects or altered compartment proportions observed following depletion of integrin pathway components (Table 3). Instead, several genes affected cap cell morphology, producing undulations similar to those observed following depletion of structural and ECM-processing components. Knockdown of *Patj* induced cap cell undulations with two independent RNAi lines, affecting 7/24 and 8/26 organs. Similar defects were observed with single RNAi lines targeting *Fak56D*, *Moe* and *l(2)gl* (Table 3). These findings suggest that maintenance of the characteristic straight morphology of cap cells depends not only on extracellular matrix organization but also on intracellular mechanisms regulating cytoskeletal organization and cell polarity.

Together, these findings reveal distinct contributions of integrin-associated proteins to LCh5 morphogenesis. Notably, across both screens, encompassing 320 RNAi lines targeting 134 genes, reproducible alterations in cap-to-ligament proportions were largely restricted to *tsp* and components of the αPS2βPS integrin-associated machinery. This striking phenotypic specificity suggests a functional relationship between Tsp-associated ECM and integrin signaling in maintaining the relative dimensions of cells during LCh5 development.

### Integrin signaling acts cell-autonomously within ligament cells to restrict elongation and permit cap cell growth

We next asked whether Tsp-dependent integrin signaling acts cell-autonomously within ligament cells to restrict their elongation and maintain their short morphology. To test this, we reduced the expression of integrin pathway components specifically in ligament cells using *repo*-Gal4 and measured cap and ligament cell lengths of each LCh5 organ (Table 5).

Ligament-specific depletion of *mys* produced pronounced ligament cell elongation accompanied by shortened cap cells, resulting in a decreased mean C/(L+SenU) ratio and 1.9- and 2.2-fold increases in the L+SenU length compared to size-matched controls, with two independent, non-overlapping RNAi lines (n = 15 and 13 hemisegments; Fig. 6 and Table 5). Knockdown of *if* produced the same phenotype with both RNAi lines tested (n = 19 and 15; L+SenU length 1.4-fold of size-matched controls for both lines; Fig. 6 and Table 5), whereas depletion of *mew* had no detectable effect (Table 5). Thus, the α-subunit specificity observed with LCh5-wide RNAi expression was recapitulated by ligament-cell-specific depletion. Notably, ligament cell-specific depletion of *if* elongated the L+SenU section in nearly every hemisegment (19/19 and 13/15), a much stronger effect than LCh5-wide depletion with *en*-Gal4 (1/14 and 6/20), suggesting that the ligament phenotype is revealed most clearly when integrin function is removed specifically from ligament cells.

**Figure 6.**
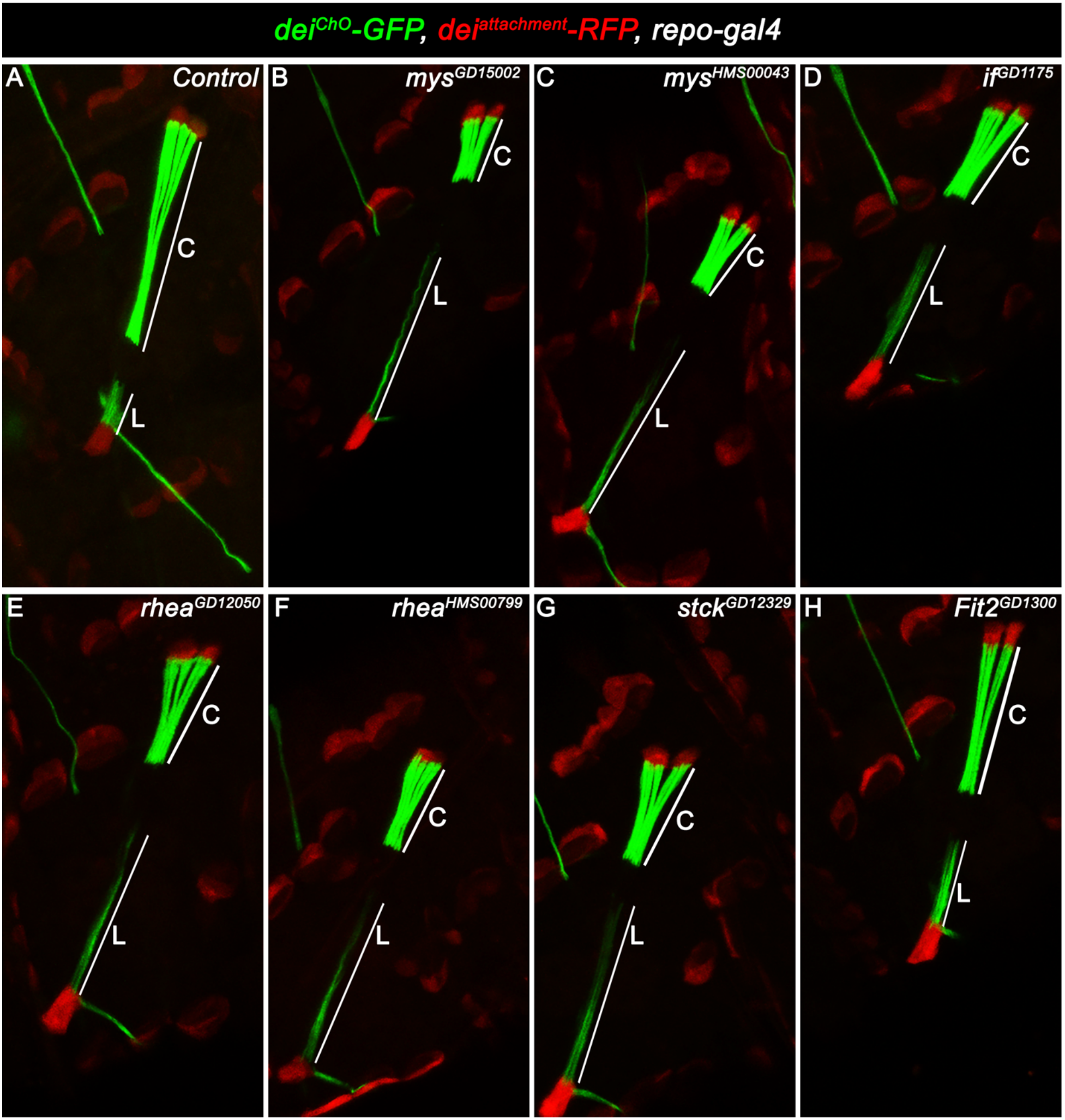
Integrin signaling in ligament cells is essential for maintaining their short morphology. (A) LCh5 organ of a 2^nd^ instar *dei^ChO^-GFP, dei^attachment^-RFP; repo-gal4* larva expressing control RNAi. (B-H) Ligament-cell depletion of *mys* (B, C), *if* (D), *rhea* (E, F), *stck* (G) and *Fit2* (H) resulted in abnormal elongation of ligament cells accompanied by reduced cap-cell length.

Depletion of intracellular integrin pathway components in ligament cells similarly reproduced the elongated-ligament/shortened-cap phenotype. Knockdown of *rhea* with two independent RNAi lines or downregulation of *stck* resulted in a 1.6- to 2.0-fold increase in L+SenU length relative to size-matched controls (Fig. 6 and Table 5). Ligament-cell-specific depletion of *Ilk* likewise produced elongated ligament cells accompanied by shortened cap cells (Fig. 6 and Table 5). Together, these results indicate that αPS2βPS integrins and their downstream effectors Talin, PINCH, and ILK are required within ligament cells to maintain the normal cap-to-ligament length balance. Notably, ligament-cell-specific depletion produced neither cap–CA detachment nor cap cell thinning across 107 hemisegments and seven genotypes, consistent with these junctional phenotypes arising from integrin requirements in a different cell type.

To further test whether the regulation of cap-to-ligament length balance was specific to integrin function within ligament cells, we depleted the same components specifically in cap cells using *β3tub^cap^*-Gal4 (Table 6). None of the eight genotypes tested produced ligament cell elongation (n=134 hemisegments; Table 6). Moreover, for every RNAi line examined with both drivers, cap-cell-specific depletion resulted in a substantially higher C/(L+SenU) ratio than ligament-cell-specific depletion of the same construct (Tables 5 and 6). Cap-cell-specific depletion occasionally produced cap cell defects, including undulation or detachment from the CA cell, but these occurred at low penetrance (Table 6). These findings demonstrate that the requirement for αPS2βPS integrin signaling in maintaining the normal cap-to-ligament length balance resides specifically within the ligament cell.

The behavior of Fit2 further distinguished the two integrin-dependent functions. Whereas *en*-Gal4-driven depletion of *Fit2* strongly disrupted cap–CA morphology without affecting the cap-to-ligament length balance, ligament-cell-specific depletion resulted in a mild reduction of the C/(L+SenU) ratio (Table 5). Although lower than in controls, this ratio was substantially higher than the ratios produced by ligament-specific depletion of *mys*, *if*, *rhea*, or *stck*, and L+SenU length was only mildly increased (1.1-fold of size-matched controls, with 7/18 LCh5 organs above the control range, versus 1.35- to 2.2-fold and nearly all hemisegments for *mys*, *if*, *rhea* and *stck*) (Fig. 6 and Table 5). Thus, Fit2 is largely dispensable within ligament cells for maintenance of the normal cell-length balance, further supporting distinct cellular requirements for integrin function in regulation of ligament cell length and cap–CA junction integrity.

Together, these findings demonstrate that the αPS2βPS–Talin–IPP integrin module acts within ligament cells to restrict their elongation. By limiting ligament cell elongation, this pathway enables normal cap cell extension, establishing ligament morphology as an actively regulated determinant of LCh5 architecture and, likely, sensory function.

### Integrin signaling in mid-embryogenesis irreversibly establishes the short morphology of ligament cells

The coupling between cap and ligament cells morphogenesis raises a temporal question: when during development must integrin signaling act to commit ligament cells to their short, non-elongating fate? To address this, we combined *repo*-Gal4-driven *mys* RNAi with temperature-sensitive Gal80 (Gal80^ts^), which represses Gal4-mediated transcription at 18°C and is inactivated at 29°C (McGuire et al., 2004) (Fig. 7). Morphology was assessed in third-instar larvae and quantified as the L+SenU length relative to total organ length, referred to here as the ligament fraction.

**Figure 7.**
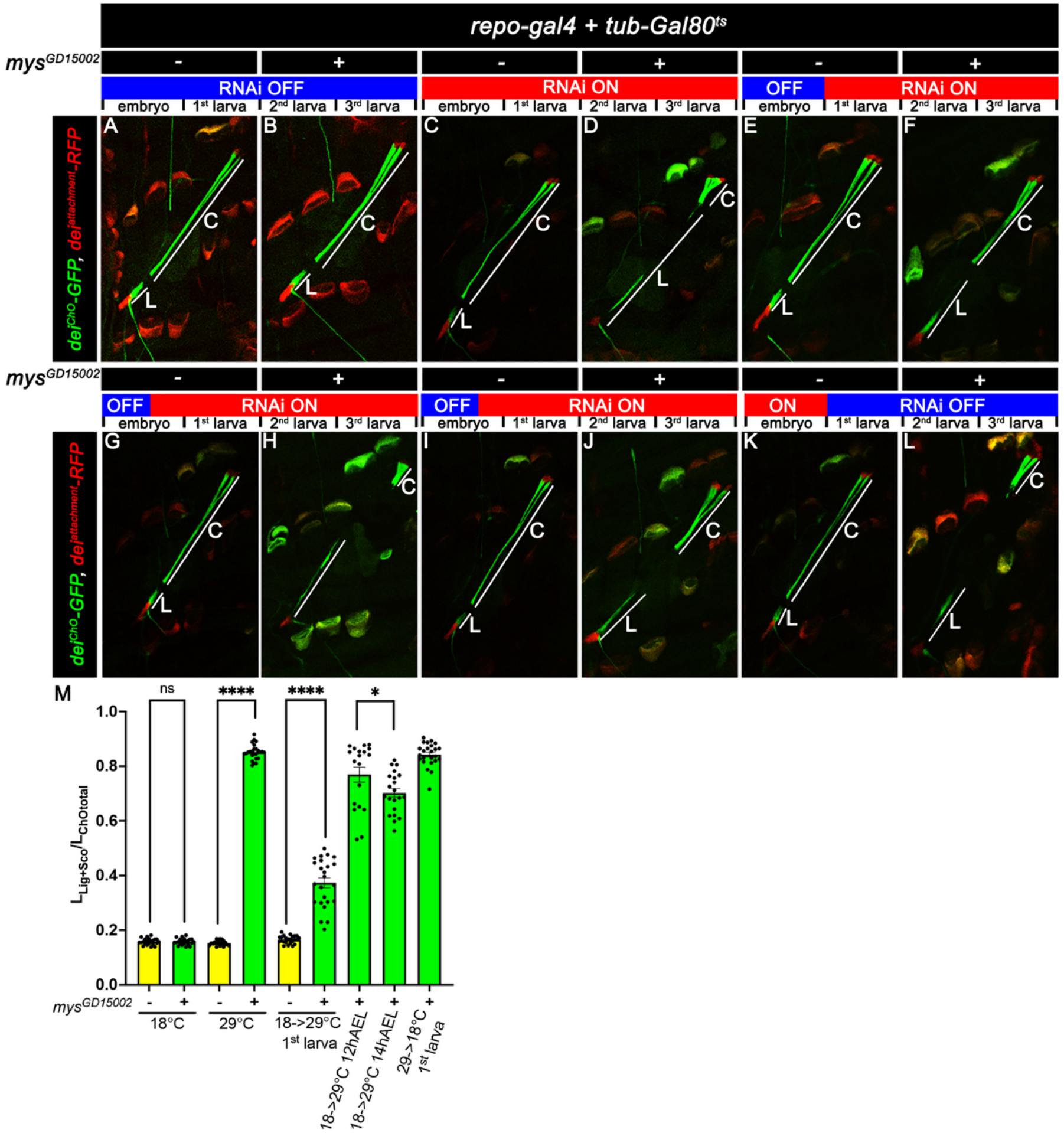
Integrin signaling during late embryogenesis restricts the ability of ligament cells to elongate during larval growth. (A–L) Representative images of LCh5 organs from larvae not expressing (−) or expressing (+) *mys RNAi* (*mys^GD15002^*). Expression of *mys^GD15002^*was temporally controlled using the temperature-sensitive Gal80 variant, Gal80^ts^, which is active at 18°C and suppresses *UAS*-driven *mys^GD15002^*expression, but inactive at 29°C, allowing *mys^GD15002^* expression. (A, B) Animals were maintained at 18°C throughout development. Under these conditions, *mys^GD15002^* expression was not induced in B. (C, D) Animals were maintained at 29°C throughout development. Continuous *mys^GD15002^* expression in D resulted in dramatic elongation of ligament (L) cells accompanied by shortening of cap (C) cells. (E, F) Animals were maintained at 18°C throughout embryonic development (*mys^GD15002^*OFF) and shifted to 29°C following 1^st^ larval hatching (*mys^GD15002^*ON). Postembryonic induction of *mys^GD15002^* resulted in only mild abnormal elongation of L cells. (G, H) Animals were maintained at 18°C (*mys^GD15002^* OFF) and shifted to 29°C at 12 hours after egg laying (hAEL; *mys^GD15002^* ON). Induction of *mys^GD15002^* at 12 hAEL resulted in an almost complete *mys RNAi* phenotype. (I, J) Animals were maintained at 18°C (*mys^GD15002^* OFF) and shifted to 29°C at 14 hAEL (*mys^GD15002^* ON). This later induction of *mys^GD15002^* resulted in a milder L cell elongation phenotype compared with induction at 12 hAEL (compare H and J). (K, L) Animals were maintained at 29°C throughout embryonic development (*mys^GD15002^*ON) and shifted to 18°C following 1^st^ instar larval hatching (*mys^GD15002^*OFF). Embryonic expression of *mys^GD15002^*, despite its postembryonic suppression, resulted in an almost complete *mys RNAi* phenotype. (M) Quantification of the experiments shown in A–L. Data are presented as means ± 95% confidence intervals (CI) and were analyzed using the Mann–Whitney U test.

Consistent with effective Gal80^ts^-mediated suppression, *repo* > *mys* RNAi animals carrying Gal80^ts^ and maintained at 18°C (RNAi off) had a ligament fraction of 0.188 ± 0.022 (n = 24), compared with 0.158 ± 0.012 in driver controls (n = 22). In contrast, the same *mys* RNAi line without Gal80^ts^ produced a ligament fraction of 0.848 ± 0.020 at 18°C (n = 24), confirming that suppression of the phenotype at this temperature was Gal80^ts^-dependent (Fig. 7 and Table 7).

Continuous *mys* depletion at 29°C produced the full phenotype, increasing the ligament fraction to 0.851 ± 0.026 (n = 25), compared with 0.153 ± 0.009 in temperature-matched controls (n = 24) (Fig. 7 and Table 7). The two compartments changed reciprocally: the L+SenU compartment was 6.6-fold longer, whereas the cap cell was reduced to 21% of the control length.

We next asked whether βPS integrin is required post-embryonically during larval growth. Embryos were maintained at 18°C (RNAi off) and shifted to 29°C (RNAi on) at first-instar hatching. These animals developed an intermediate phenotype, with a ligament fraction of 0.373 ± 0.089 (n = 24), compared with 0.166 ± 0.014 in shift-matched controls (n = 24) (Fig. 7 and Table 7). Thus, *mys* depletion initiated at hatching was sufficient to alter relative cell lengths but produced a substantially weaker phenotype than continuous depletion, indicating a major requirement for βPS integrin function before hatching.

To further narrow the period of integrin requirement, *mys* RNAi was induced at different times during embryogenesis. Shifting embryos to 29°C at 14 h after egg laying (AEL) resulted in a ligament fraction of 0.703 ± 0.076 (n = 21), whereas shifting at 12 h AEL increased the ligament fraction to 0.786 ± 0.098 (n = 19) (Fig. 7 and Table 7). The two conditions differed significantly, and the phenotype following the 12 h AEL shift was not significantly different from that produced by continuous knockdown. Thus, initiating *mys* RNAi by 12 h AEL was sufficient to produce a phenotype resembling that of continuous depletion, whereas induction at 14 h AEL produced a significantly weaker effect, narrowing the critical window of βPS depletion to mid-embryogenesis.

The commitment established during this period is irreversible. Animals maintained at 29°C throughout embryogenesis and shifted to 18°C at hatching (restoring integrin expression in ligament cells throughout larval life) had a ligament fraction of 0.843 ± 0.043 (n = 24), indistinguishable from that produced by continuous knockdown (Fig. 7 and Table 7). Thus, restoring conditions permissive for integrin expression at hatching did not rescue the altered cap and ligament proportions. These findings indicate that βPS integrin is required during embryogenesis for establishing normal LCh5 proportions, a requirement that cannot be compensated for by postembryonic βPS activity. These findings also suggest that embryonic integrin signaling does not act continuously to maintain ligament cell morphology but triggers a downstream program that becomes self-sustaining during larval life.

Consistent with an embryonic requirement, altered cap and ligament proportions were already detectable at hatching. In first-instar larvae, *repo* > *mys* RNAi animals had a higher ligament fraction than controls (0.605 ± 0.043 versus 0.428 ± 0.043; n = 20 each), while total labeled organ length was unchanged (Fig. 7 and Table 7). This difference became more pronounced during larval development. In control larvae, cap cells elongated more extensively than ligament cells during larval growth. Consequently, the proportion of total organ length occupied by the L+SenU section decreased from 0.428 in 1^st^ instar larvae to 0.153 in 3^rd^ instar larvae. In *mys* RNAi animals, the opposite occurred: the L+SenU region became increasingly dominant and increased from 0.605 to 0.851. Thus, βPS depletion alters the relative dimensions of the cap and ligament compartments by hatching, and these differences are amplified during subsequent larval growth.

Together, these experiments define the molecular and temporal basis of ligament cell morphology. βPS integrin-mediated signaling acts cell-autonomously within ligament cells during a defined window in embryogenesis to irreversibly establish their short, non-elongated morphology, creating the structural framework for subsequent cap cell elongation and mechanical maturation.

## Discussion

Despite their common developmental origin and mechanical interconnection, cap and ligament cells respond very differently to the stretching imposed by larval growth. Understanding how lineage-related cells acquire and maintain distinct morphologies and mechanical properties in response to forces experienced during development is a fundamental question in developmental biology. Our findings revealed ECM-associated mechanisms that contribute to both the structural integrity of the organ and the regulation of its characteristic cell-length proportions. Specifically, we demonstrate that Tsp and αPS2βPS integrin-associated machinery are required to maintain the normal balance between cap- and ligament-cell elongation by restricting ligament-cell elongation. This function is at least partially separable from the role of integrins in attachment stability and depends primarily on their activity within the ligament cells. Interestingly, the predominantly embryonic requirement for integrin signaling suggests that the mechanisms controlling differential elongation may be established before the organ becomes functional and undergoes developmental stretching.

Our screen identified several ECM components required to maintain the characteristic straight morphology of cap cells, including laminins and Multiplexin, a collagen XV/XVIII family member. Although the basement-membrane components whose localization was examined in this study are broadly distributed, within LCh5 they enwrap the entire organ and colocalize with Pericardin (Prc), which has a more restricted distribution. We previously demonstrated that Prc affects the mechanical properties of the cap cell ECM and is required for normal cap-cell mechanical properties and mechanosensory function (Hassan et al., 2019). These localization patterns, together with the cap-cell undulation observed upon depletion of these ECM components, support a model in which the surrounding ECM forms a sleeve-like scaffold that provides structural support to these slender cells. Such a scaffold may stabilize cap cells during their extensive post-embryonic elongation while maintaining their ability to transmit muscle contraction-induced mechanical forces to the sensory neuron. The requirement for ECM-modifying enzymes, including Tld metalloproteases and lysyl oxidases, further suggests that proper matrix assembly and potentially dynamic remodeling are important for maintaining this structural support during larval growth. Whether these other ECM molecules contribute to cap-cell mechanical properties, as has been demonstrated for Prc, remains to be established.

In addition to providing structural support to elongated cap cells, the ECM must ensure stable anchoring of the LCh5 organ to the cuticle at both ends. The continuous enlargement of these attachment sites during larval growth likely helps maintain stable adhesion as the organ increases in size. Our screen identified two junctional ECM proteins, Tsp and Tig, and the αPS2βPS integrin complex as important regulators of LCh5 attachment stability. Interestingly, these proteins are also involved in stabilizing *Drosophila* muscle attachment sites (Bunch et al., 1998; Chanana et al., 2007; Fogerty et al., 1994; Subramanian et al., 2007; Vanhoutte et al., 2016), which share developmental and molecular similarities with ChO attachments (Hasson et al., 2017). However, the timing of Tsp accumulation differs between the two systems. At myotendinous junctions (MTJs), Tsp is expressed in tendon precursors during embryonic stages 12–13 and becomes concentrated at the junctions by stage 16 (Hasson et al., 2017). In contrast, Tsp becomes detectable at ChO attachment sites during late embryogenesis (stage 17), initially around the LA cell and at much lower levels at the cap–CA junction. Strong enrichment at both junctions occurs during larval development (Greenblatt Ben-El et al., 2017). This delayed accumulation coincides with extensive organ growth and increasing mechanical loads, raising the possibility that these forces promote the maturation and reinforcement of integrin-mediated adhesions, accompanied by increased Tsp incorporation into the junctional ECM.

Our findings further extend the known role of Tsp in ChO attachment by demonstrating its requirement for maintaining the ligament–LA junction. Notably, previous lineage-restricted depletion of Tsp affected only the cap–CA junction (Greenblatt Ben-El et al., 2017), whereas ubiquitous depletion also caused ligament–LA detachment. This difference raises the possibility that Tsp required for ligament–LA attachment is supplied by LA cells, consistent with the previously reported expression of *tsp* in these cells during late embryogenesis (Greenblatt Ben-El et al., 2017). Interestingly, the two attachment sites responded differently to integrin depletion. Consistent with the previously work, βPS depletion caused defects at the cap-CA junction (Greenblatt Ben-El et al., 2017; Hassan et al., 2018) but did not result in ligament–LA detachment. This difference raises the possibility that additional adhesion mechanisms, potentially involving other ECM receptors, compensate for reduced βPS integrin activity at the ligament–LA junction but not at the cap–CA junction.

Beyond their role in maintaining attachment stability, our screen revealed an unexpected role for Tsp and the αPS2βPS integrin in regulating the relative elongation of cap and ligament cells. Depletion of either *tsp* or components of the αPS2βPS integrin complex disrupted the cap-to-ligament length balance. Given the established interaction between Tsp and αPS2βPS integrins at MTJs (Subramanian et al., 2007), these similar phenotypes raise the possibility that a related ligand–receptor interaction contributes to the regulation of cell elongation in ChOs. Importantly, integrin depletion caused pronounced ligament-cell elongation without disruption of the ligament–LA attachment, indicating that the length phenotype does not necessarily result from junctional failure. Nevertheless, the absence of detachment does not exclude changes in adhesion strength that could influence ligament-cell elongation.

Cell-type-specific experiments further demonstrated that αPS2βPS integrin and its associated intracellular machinery are required within ligament cells to maintain the normal cap-to-ligament length balance during larval growth. Despite their connection in series, cap and ligament cells respond differently to developmental stretching (Halachmi et al., 2016). Interestingly, integrin depletion resulted in excessive ligament-cell elongation accompanied by reduced cap-cell elongation, without affecting the total organ length. These reciprocal changes suggest that integrin-associated mechanisms contribute to the coordination of elongation between the two cell types. One possibility is that integrin signaling helps ligament cells resist developmental stretching, thereby allowing cap cells to accommodate most of the organ’s elongation. Alternatively, integrins may regulate ligament-cell growth or mechanical coupling to the surrounding ECM. Although the distinct mechanical behaviors of cap and ligament cells favor the first possibility, whether integrin depletion directly alters ligament-cell mechanical properties remains to be established.

A possible link between integrin signaling and ligament-cell elongation is the organization of the microtubule cytoskeleton. Previous work demonstrated that specific tubulin isoforms and microtubule-associated proteins contribute to maintaining the differential elongation of ChO cells, with ligament cells particularly sensitive to depletion of αTub85E (Hassan et al., 2018). Interestingly, the spectraplakin Shot, which links actin filaments and microtubules (Lee and Kolodziej, 2002), contributes to the organization of microtubule arrays required for mechanical stability in tendon cells (Subramanian et al., 2003). Shot also localizes to ChO attachment sites, and its depletion affects CA and ligament-cell elongation (Hassan et al., 2018). These observations raise the possibility that integrin-associated machinery, including Talin and the IPP complex, regulates ligament-cell elongation by influencing microtubule organization, potentially through the actin–microtubule crosslinking protein Shot. Such interactions could regulate microtubule organization and the mechanical response of ligament cells to developmental stretching. However, a functional connection between integrin signaling and microtubule organization in ligament cells remains to be demonstrated. The absence of pronounced length phenotypes following depletion of several actin-associated proteins does not exclude their involvement, as redundancy or incomplete knockdown may have masked individual requirements.

The potential relationship between integrin-associated machinery and cytoskeletal organization raises the question of how these mechanisms are stablished during development. Previous work demonstrated that depletion of the transcription factors Sr and Dei causes the abnormal ligament-cell elongation phenotype as depletion of integrin does (Hassan et al., 2018). Sr is required for ligament-cell differentiation and regulates several components involved in cytoskeletal organization and adhesion, including αTub85E, Shot, and βPS integrin (Klein et al., 2010). Dei, although dispensable for αTub85E expression in ligament cells, has been implicated in regulating βPS integrin expression and cell adhesion in other tissues (Egoz-Matia et al., 2011; Hassan et al., 2018). These findings raise the possibility that Sr and Dei regulate distinct but partially overlapping mechanisms that establish the characteristic short morphology of ligament cells. Integrin-associated machinery may represent one link between these transcriptional programs and the cellular interactions with the ECM required for normal cell elongation; whether they regulate this machinery in ligament cells, directly or indirectly, remains to be determined.

Our temporal experiments suggest that integrin activity is particularly important during embryogenesis for establishing normal cap-to-ligament length proportions. Abnormal proportions were already apparent in first-instar larvae and persisted when RNA interference was relieved after hatching, whereas knockdown initiated at hatching produced a weaker phenotype. These findings raise the possibility that integrin-associated machinery contributes to establishing ligament-cell properties during embryogenesis, before extensive larval growth. Interestingly, recent work demonstrated that Prc is required during a critical developmental period for normal cap-cell mechanics and that reducing muscle-generated forces during embryogenesis causes persistent defects in ChO mechanosensory function (Hassan et al., 2026). Thus, early interactions between cells, the ECM, and mechanical forces may influence the subsequent mechanical identity of ChO cells.

## Materials and Methods

### Fly strains

The following fly strains were used in the study: *Luc^JF01355^*RNAi line (#31603, Bloomington Drosophila Stock Center (BDSC)) was used as a control RNAi with all driver lines. *vkg^fTRG00595.sfGFP-TVPTBF^* (v318167, Vienna Drosophila Resource Center (VDRC)), *LanA^fTRG00574.sfGFP-TVPTBF^* (v318155, VDRC), *LanB1^fTRG00681.sfGFP-TVPTBF^* (v318180, VDRC), *dei^ChO^-GFP, dei^attachment^-RFP, en-gal4* (Hassan et al., 2018), *tub-Gal80^ts^/TM2* (#7017, BDSC). The lines *dei^ChO^-GFP, dei^attachment^-RFP; repo-gal4/TM6, AbdA-lacZ and mys^GD15002^/CyO, Kr>GFP; tub-Gal80^ts^/ TM6, AbdA-lacZ* were generated by standard genetic crosses. *dei^attachment^-RFP; dei^ChO^-GFP, da-gal4/TM6, AbdA-lacZ* was generated by meiotic recombination following standard genetic crosses. The full list of RNAi lines used in this study is provided in the Supplementary Table 2.

### Phenotypic analysis in 2nd instar larvae

Virgin females from the appropriate driver lines were crossed to *UAS-RNAi* males and allowed to lay eggs for 24 h at 29°C. The parents were then discarded, and the progeny were collected 24 h later. The larvae were washed and fixed overnight in 4% formaldehyde in PBS. Following fixation, the larvae were washed twice with PBT and once with PBS. The larvae were then stored in mounting medium (DAKO, DakoCytomation, Denmark) at 4°C and mounted and imaged within one week.

### Temporal *mys* knockdown experiments

Virgin females of the *dei^ChO^-GFP, dei^attachment^-RFP; repo-gal4/TM6, AbdA-lacZ* genotype were crossed to *mys^GD15002^/CyO, Kr>GFP; tub-Gal80^ts^/ TM6, AbdA-lacZ* males and allowed to lay eggs for 2h. The parents were then transferred to fresh vials, and the vials containing the eggs were incubated at either 18°C or 29°C according to the experimental temperature regimen. At the appropriate developmental stage, defined by hours after egg laying (hAEL), the progeny were shifted from 18°C to 29°C or vice versa, depending on the experimental conditions. The larvae were examined at the wandering 3^rd^ instar stage.

### Quantification and statistical analysis

Cap cell undulation, cap cell thinning, cap–CA detachment and ligament–LA detachment were scored per abdominal hemisegment and phenotype frequencies are given as affected hemisegments over all scored hemisegments. Each genotype was compared with control larvae expressing *Luc RNAi* (JF01355) under the same Gal4 driver. Phenotype frequencies were compared with the control using a one-sided Fisher’s exact test (alternative hypothesis: a higher frequency than in controls). An RNAi line was scored as positive when p < 0.05 and the phenotype frequency was at least 15%, and a gene was considered a hit when at least two independent, non-overlapping RNAi lines were positive.

For length measurements, cap cell length and total GFP-labelled organ length were measured in each hemisegment, and L+SenU length was calculated as total minus cap length. Size-specific limits were derived from the same-driver control hemisegments by regressing log(C/(L+SenU)) on log(total organ length) and L+SenU length on total organ length; a hemisegment was scored as having a reduced ratio when it fell below the 2.5th percentile of the residual distribution (mean − 1.96 SD) and an elongated ligament when L+SenU length exceeded the 97.5th percentile. The proportions of hemisegments beyond these limits were compared with controls by one-sided Fisher’s exact tests, using the same 15% and two-line criteria. Hemisegments in which ligament cells appeared detached from the LA cell were excluded from length analyses and reported separately.

Associations between phenotypes within hemisegments were tested with two-sided Fisher’s exact tests, and C/(L+SenU) ratios of the same RNAi line under different drivers were compared with two-sided Mann–Whitney tests. GraphPad Prism 10 was used for statistical analyses (GraphPad Prism version 10, GraphPad Software, Boston, Massachusetts USA, www.graphpad.com).

