## Supplementary material for "Embryonic integrin signaling establishes a program of differential cell elongation in a growing proprioceptive organ": Tables 1-6

### Table 1. Ubiquitous (*da*-Gal4) knockdown of ECM and ECM-associated genes

| Gene | RNAi line (BDSC or VDRC stock #) | Hemi-segments | Undulated cap | Cap–CA detachment | Cap thinning | Reduced cap/ligament ratio | Ligament–LA detachment | Positive |
|---|---|---|---|---|---|---|---|---|
| ***Control*** | | | | | | | | |
| *Luc* | 31603 | 42 | 2 (5%) | 0 | 0 | — | 0 | |
| ***Validation*** | | | | | | | | |
| *prc* | 100357 | 61 | 32 (52%) | 0 | 0 | 0 | 0 | U |
| *prc* | 41320 | 44 | 26 (59%) | 0 | 0 | 0 | 0 | U |
| ***Screen*** | | | | | | | | |
| *AdamTS-A* | 110157 | 41 | 0 | 0 | 0 | 0 | — | |
| *AdamTS-A* | 110157 | 38 | 0 | 0 | 0 | 0 | — | |
| *atk* | 31044 | 24 | 6 (25%) | 0 | 0 | 0 | — | U |
| *atk* | 64917 | 21 | 9 (43%) | 1 (5%) | 0 | 0 | — | U |
| *atk* | 100110 | 30 | 1 (3%) | 0 | 0 | 0 | — | |
| *Cad86C* | 21327 | 49 | 0 | 0 | 0 | 0 | — | |
| *Cad86C* | 26587 | 97 | 0 | 0 | 0 | 0 | — | |
| *Cad86C* | 109690 | 20 | 0 | 0 | 0 | 0 | — | |
| *Cad86C* | 109690 | 42 | 0 | 0 | 0 | 0 | — | |
| *Cad87A* | 8578 | 28 | 0 | 0 | 0 | 0 | — | |
| *Cad87A* | 105901 | 63 | 1 (2%) | 0 | 0 | 0 | — | |
| *Cad99C* | 27212 | 42 | 0 | 0 | 0 | 0 | — | |
| *Cad99C* | 27212 | 13 | 0 | 0 | 0 | 0 | — | |
| *Cad99C* | 27216 | 49 | 0 | 0 | 0 | 0 | — | |
| *Cad99C* | 27216 | 49 | 0 | 1 (2%) | 0 | 0 | — | |
| *Cad99C* | 27216 | 62 | 0 | 3 (5%) | 0 | 0 | — | |
| *Cad99C* | 27216 | 64 | 0 | 0 | 0 | 0 | — | |
| *CadN2* | 47538 | 49 | 2 (4%) | 1 (2%) | 1 (2%) | 0 | — | |
| *CadN2* | 47538 | 14 | 0 | 1 (7%) | 0 | 0 | — | |
| *cals* | 37291 | 47 | 0 | 0 | 0 | 0 | — | |
| *cals* | 105111 | 49 | 0 | 0 | 0 | 0 | — | |
| *CG5550* | 31001 | 40 | 0 | 1 (2%) | 0 | 0 | — | |
| *CG5550* | 31001 | 42 | 0 | 0 | 0 | 0 | — | |
| *clos* | 32122 | 91 | 0 | 0 | 0 | 0 | — | |
| *clos* | 104142 | 71 | 0 | 0 | 0 | 0 | — | |
| *clos* | 108711 | 18 | 0 | 0 | 0 | 0 | — | |
| *Col4a1 (Cg25c)* | 44520 | 27 | 3 (11%) | 1 (4%) | 0 | 0 | — | |
| *Col4a1 (Cg25c)* | 44520 | 29 | 3 (10%) | 1 (3%) | 0 | 0 | — | |
| *Col4a1 (Cg25c)* | 104536 | — | — | — | — | — | — | not scored |
| *Ddr* | 29720 | 77 | 0 | 0 | 0 | 0 | — | |

| Gene | RNAi line (BDSC or VDRC stock #) | Hemi-segments | Undulated cap | Cap–CA detachment | Cap thinning | Reduced cap/ligament ratio | Ligament–LA detachment | Positive |
|---|---|---|---|---|---|---|---|---|
| *Ddr* | 51719 | 48 | 0 | 0 | 0 | 0 | — | |
| *Dg* | 34895 | 18 | 0 | 1 (6%) | 0 | 0 | — | |
| *dpy* | 7836 | 84 | 0 | 1 (1%) | 0 | 0 | — | |
| *ds* | 4313 | 42 | 0 | 0 | 0 | 0 | — | |
| *eys* | 22541 | 74 | 1 (1%) | 0 | 0 | 0 | — | |
| *eys* | 31513 | 31 | 0 | 0 | 0 | 0 | — | |
| *eys* | 31514 | 27 | 2 (7%) | 0 | 0 | 0 | — | |
| *eys* | 32599 | 64 | 1 (2%) | 0 | 0 | 0 | — | |
| *eys* | 33764 | 14 | 0 | 0 | 0 | 0 | — | |
| *eys* | 33766 | 20 | 5 (25%) | 0 | 0 | 0 | — | U |
| *eys* | 106892 | 62 | 0 | 0 | 0 | 0 | — | |
| *ft* | 9396 | 84 | 0 | 0 | 0 | 0 | — | |
| *Gfrl* | 36443 | 77 | 0 | 0 | 0 | 0 | — | |
| *Gfrl* | 103213 | 47 | 0 | 1 (2%) | 0 | 0 | — | |
| *Gfrl* | 103523 | 100 | 0 | 1 (1%) | 0 | 0 | — | |
| *Hml* | 37005 | 30 | 5 (17%) | 0 | 0 | 1 (3%) | — | |
| *Hml* | 37006 | 26 | 1 (4%) | 0 | 0 | 0 | — | |
| *Hml* | 50640 | 22 | 3 (14%) | 0 | 0 | 0 | — | |
| *Hml* | 108935 | 22 | 3 (14%) | 0 | 0 | 0 | — | |
| *ItgapS5* | 6647 | 70 | 0 | 0 | 0 | 0 | — | |
| *ItgapS5* | 100120 | 61 | 0 | 0 | 0 | 0 | — | |
| *kirre* | 27227 | 42 | 1 (2%) | 0 | 0 | 0 | — | |
| *kirre* | 109585 | 56 | 0 | 1 (2%) | 0 | 0 | — | |
| *kug* | 27114 | 76 | 0 | 0 | 0 | 0 | — | |
| *kug* | 110390 | 49 | 0 | 0 | 0 | 0 | — | |
| *LanA* | — | — | — | — | — | — | — | not scored |
| *LanB1* | 42616 | 22 | 4 (18%) | 0 | 0 | 0 | — | |
| *LanB2* | 42559 | 14 | 0 | 0 | 0 | 0 | — | |
| *LanB2* | 42560 | 23 | 6 (26%) | 0 | 0 | 0 | — | U |
| *LanB2* | 62002 | 22 | 5 (23%) | 1 (5%) | 0 | 0 | — | U |
| *LanB2* | 104013 | 12 | 5 (42%) | 0 | 0 | 0 | — | U |
| *Lcp1* | 106030 | 39 | 0 | 0 | 0 | 0 | — | |
| *Lcp2* | 12537 | 103 | 0 | 2 (2%) | 0 | 0 | — | |
| *Lcp3* | 107682 | 78 | 2 (3%) | 0 | 0 | 0 | — | |
| *Loh* | 31020 | 48 | 0 | 1 (2%) | 0 | 0 | — | |
| *Loh* | 31021 | 53 | 4 (8%) | 0 | 0 | 0 | — | |
| *Loxl1* | 107435, 16619 | 24 | 4 (17%) | 0 | 0 | 0 | — | |

| Gene | RNAi line (BDSC or VDRC stock #) | Hemi-segments | Undulated cap | Cap–CA detachment | Cap thinning | Reduced cap/ligament ratio | Ligament–LA detachment | Positive |
|---|---|---|---|---|---|---|---|---|
| *Loxl1* | 17259, 16619 | 23 | 6 (26%) | 0 | 0 | 0 | — | U |
| *Mcr* | 65896 | 16 | 0 | 4 (25%) | 0 | 0 | — | D |
| *mfas* | 103621 | 40 | 0 | 0 | 0 | 0 | — | |
| *MMP2* | 65395 | 24 | 3 (12%) | 0 | 0 | 0 | — | |
| *Mp* | 38188 | 30 | 7 (23%) | 0 | 0 | 0 | — | U |
| *Mp* | 38189 | 27 | 0 | 0 | 0 | 0 | — | |
| *Mp* | 52981 | 22 | 1 (5%) | 0 | 0 | 0 | — | |
| *Mp* | 100846 | 29 | 4 (14%) | 0 | 0 | 0 | — | |
| *Mp* | 107319 | 21 | 7 (33%) | 0 | 0 | 0 | — | U |
| *mspo* | 107608 | 48 | 0 | 0 | 0 | 0 | — | |
| *Muc12Ea* | 50435 | 119 | 0 | 0 | 0 | 0 | — | |
| *Muc12Ea* | 100177 | 28 | 0 | 1 (4%) | 0 | 0 | — | |
| *Muc18B* | 100171 | 35 | 0 | 0 | 0 | 0 | — | |
| *Muc26B* | 104205 | 68 | 0 | 3 (4%) | 0 | 0 | — | |
| *Muc26B* | 104205 | 49 | 0 | 0 | 0 | 0 | — | |
| *Muc4B* | 42100 | 24 | 0 | 0 | 0 | 0 | — | |
| *Muc4B* | 42101 | 24 | 1 (4%) | 0 | 0 | 0 | — | |
| *Muc4B* | 67940 | 22 | 5 (23%) | 0 | 0 | 0 | — | U |
| *Muc4B* | 76072 | 23 | 6 (26%) | 0 | 0 | 0 | — | U |
| *Muc55B* | 6425 | 42 | 0 | 0 | 0 | 0 | — | |
| *Muc55B* | 104747 | 40 | 0 | 0 | 0 | 0 | — | |
| *Muc68Ca* | 48500 | 112 | 4 (4%) | 0 | 0 | 0 | — | |
| *Muc68D* | 105461 | 49 | 0 | 0 | 0 | 0 | — | |
| *Muc68E* | 48553 | 61 | 0 | 0 | 0 | 0 | — | |
| *Muc82C* | 100030 | 97 | 0 | 0 | 0 | 0 | — | |
| *Muc86D* | 25489 | 42 | 0 | 0 | 0 | 0 | — | |
| *Muc91C* | 9867 | 27 | 0 | 0 | 0 | 0 | — | |
| *Mur11Da* | 100061 | 103 | 0 | 0 | 0 | 0 | — | |
| *Mur29B* | 46704 | 39 | 0 | 0 | 0 | 0 | — | |
| *Mur89F* | 100128 | 35 | 0 | 0 | 0 | 0 | — | |
| *Ndg* | 13280 | 24 | 7 (29%) | 0 | 0 | 0 | — | U |
| *Ndg* | 62902 | 29 | 0 | 1 (3%) | 0 | 0 | — | |
| *Ndg* | 109625 | 26 | 1 (4%) | 0 | 0 | 0 | — | |
| *nolo (CG31619)* | 33102 | 49 | 0 | 0 | 0 | 1 (2%) | — | |
| *nolo (CG31619)* | 104736 | 28 | 0 | 0 | 0 | 0 | — | |
| *obst-A* | 12935 | 35 | 0 | 1 (3%) | 0 | 0 | — | |
| *Plod* | 45484 | 7 | 0 | 0 | 0 | 0 | — | |

| Gene | RNAi line (BDSC or VDRC stock #) | Hemi-segments | Undulated cap | Cap–CA detachment | Cap thinning | Reduced cap/ligament ratio | Ligament–LA detachment | Positive |
|---|---|---|---|---|---|---|---|---|
| *Ppn* | 16523 | 42 | 0 | 0 | 0 | 0 | — | |
| *pyd* | 104159 | 21 | 0 | 0 | 0 | 0 | — | |
| *pyd* | 104159 | 56 | 0 | 0 | 0 | 0 | — | |
| *Sog* | 37405 | 24 | 2 (8%) | 0 | 1 (4%) | 0 | — | |
| *Sog* | 37407 | 20 | 1 (5%) | 0 | 0 | 0 | — | |
| *Sog* | 67975 | 26 | 7 (27%) | 1 (4%) | 0 | 0 | — | U |
| *Sog* | 105853 | 20 | 3 (15%) | 0 | 0 | 0 | — | |
| *SPARC* | 16677 | 23 | 2 (9%) | 0 | 0 | 0 | — | |
| *SPARC* | 16678 | 25 | 2 (8%) | 0 | 0 | 0 | — | |
| *SPARC* | 40885 | 25 | 0 | 2 (8%) | 0 | 0 | — | |
| *SPARC* | 100566 | 20 | 1 (5%) | 0 | 0 | 0 | — | |
| *Tace* | 2733 | 35 | 0 | 0 | 0 | 0 | — | |
| *Tig* | 28255 | 27 | 1 (4%) | 2 (7%) | 0 | 0 | — | |
| *Tig* | 28257 | 25 | 2 (8%) | 1 (4%) | 11 (44%) | 0 | — | T |
| *Tig* | 31570 | 29 | 6 (21%) | 3 (10%) | 0 | 0 | — | U |
| *Tig* | 100036 | 27 | 0 | 0 | 7 (26%) | 0 | — | T |
| *Timp* | 15372 | 28 | 0 | 1 (4%) | 0 | 0 | — | |
| *Timp* | 109427 | 49 | 0 | 0 | 0 | 0 | — | |
| *tld* | 1215 | 25 | 6 (24%) | 0 | 0 | 0 | — | U |
| *tld* | 1216 | 22 | 7 (32%) | 0 | 0 | 0 | — | U |
| *tld* | 51507 | 23 | 9 (39%) | 0 | 0 | 2 (9%) | — | U |
| *tld* | 100930 | 27 | 3 (11%) | 0 | 0 | 0 | — | |
| *tld + tok* | 100930, 2656 | 19 | 1 (5%) | 0 | 3 (16%) | 0 | — | T |
| *tld + tok* | 1215, 2656 | 21 | 3 (14%) | 0 | 1 (5%) | 0 | — | |
| *Tok* | 2656 | 27 | 5 (19%) | 1 (4%) | 1 (4%) | 0 | — | |
| *Tok* | 66320 | 18 | 5 (28%) | 0 | 0 | 0 | — | U |
| *Tok* | 110432 | 24 | 5 (21%) | 0 | 0 | 0 | — | |
| *trol* | 22642 | 25 | 4 (16%) | 0 | 0 | 0 | — | |
| *trol* | 24549 | 27 | 2 (7%) | 0 | 0 | 0 | — | |
| *trol* | 38298 | 15 | 1 (7%) | 1 (7%) | 0 | 0 | — | |
| *trol* | 38298 | 20 | 1 (5%) | 0 | 0 | 1 (5%) | — | |
| *trol* | 110494 | 29 | 0 | 0 | 0 | 0 | — | |
| *tsp* ¶ | 7535 | 42 | 0 | 7 (17%) | 0 | 20/35 (57%)‡ | 7 (17%) | D,L,R |
| *tsp* ¶ | 100721 | 70 | 0 | 13 (19%) | 0 | 7/49 (14%)‡ | 15 (21%) | D,L |
| *TwdlB* | 103294 | 55 | 0 | 0 | 0 | 0 | — | |
| *TwdlE* | 24867 | 60 | 0 | 0 | 0 | 0 | — | |
| *TwdlE* | 107483 | 21 | 0 | 0 | 0 | 0 | — | |
| *TwdlG* | 15944 | 42 | 0 | 0 | 0 | 0 | — | |

| Gene | RNAi line (BDSC or VDRC stock #) | Hemi-segments | Undulated cap | Cap–CA detachment | Cap thinning | Reduced cap/ligament ratio | Ligament–LA detachment | Positive |
|---|---|---|---|---|---|---|---|---|
| *TwdlG* | 100335 | 47 | 0 | 1 (2%) | 0 | 0 | — | |
| *TwdlT* | 41335 | 42 | 0 | 0 | 0 | 0 | — | |
| *TwdlT* | 107928 | 69 | 0 | 0 | 0 | 0 | — | |
| *TwdlV* | 102989 | 75 | 0 | 0 | 0 | 0 | — | |

Counts are affected hemisegments (% of hemisegments scored in that experiment). Repeated experiments with the same RNAi line are listed separately in consecutive rows and were not pooled.

Reduced cap/ligament ratio: hemisegments with a visibly reduced ratio of cap to ligament length, scored by eye; for *tsp* (‡) the value is the number of hemisegments below the size-specific threshold from measurements (Table 2). Ligament–LA detachment was observed only with *tsp* knockdown.

**Positive:** phenotypes for which the line met the criterion, a frequency of at least 15% that is higher than in the *da* > Luc RNAi control (one-sided Fisher's exact test, $p < 0.05$). U, cap undulation; D, cap–CA detachment; T, cap thinning; L, ligament–LA detachment; R, reduced size-adjusted cap/ligament ratio. Genes with at least two positive lines for the same phenotype: undulation, *LanB2*, *Mp*, *Muc4B*, *tld* and *atk*; cap thinning, *Tig*; cap–CA and ligament–LA detachment, *tsp*.

*prc* is shown as a validation control and is excluded from the totals.

¶ (after the gene name) *tsp* data come from Table 2.

**Totals (excluding *prc*): 132 RNAi lines or line combinations, 67 genes, 144 experiments and 5,854 hemisegments.**

### Table 2. Cap and ligament dimensions after tsp knockdown

| Gene | RNAi line (BDSC or VDRC stock #) | Larvae | Hemi-segments | Organ length (µm) | Cap length (µm) | L+SenU length (µm) | C/(L+SenU) | Below 1.1 | Reduced ratio (size-matched) | Elongated ligament (size-matched) | L+SenU, fold of size-matched control | Positive |
|---|---|---|---|---|---|---|---|---|---|---|---|---|
| *Luc (control)* | 31603 | 7 | 42 | 108.1 ± 27.1 | 65.0 ± 21.3 | 43.1 ± 7.0 | 1.49 ± 0.35 | 6/42 (14%) | 0/42 (0%) | 0/42 (0%) | 1.00 | |
| *tsp* | 7535 | 7 | 35 | 100.7 ± 13.2 | 51.5 ± 10.2 | 49.2 ± 5.4 | 1.05 ± 0.20 | 20/35 (57%) | 20/35 (57%) | 16/35 (46%) | 1.19 | R,E |
| *tsp* | 100721 | 11 | 49 | 92.0 ± 20.7 | 50.5 ± 16.5 | 41.5 ± 6.5 | 1.21 ± 0.31 | 24/49 (49%) | 7/49 (14%) | 5/49 (10%) | 1.05 | |

Lengths are mean ± SD (µm). Organ length = total GFP-labelled length; L+SenU = organ length − cap length. Hemisegments with ligament–LA detachment are excluded (7 for line 7535 and 13 for line 100721).

**Below 1.1:** C/(L+SenU) < 1.1, given for reference. **Reduced ratio** and **elongated ligament:** hemisegments beyond the size-specific limits derived from the *da* > Luc RNAi control (Methods). **L+SenU fold:** mean observed length divided by the mean expected for control organs of the same total length.

**Positive:** R, reduced ratio; E, elongated ligament (at least 15% and one-sided Fisher's exact test $p < 0.05$ versus control).

**Table 3. LCh5-directed (*en*-Gal4) knockdown of ECM receptors, adhesion and cytoskeletal genes**

| Gene | RNAi line (BDSC or VDRC stock #) | Hemi-segments | Undulated cap | Cap–CA detachment | Cap thinning | Reduced cap/ligament ratio | Positive |
|---|---|---|---|---|---|---|---|
| ***Control*** | | | | | | | |
| *Luc* | 31603 | 47 | 3 (6%) | 2 (4%) | 0 | — | |
| ***Validation*** | | | | | | | |
| *prc* | 41320 | 42 | 0 | 0 | 0 | 0 | |
| *prc* | 100357 | 42 | 0 | 0 | 0 | 0 | |
| ***Screen*** | | | | | | | |
| *Act42A* | 12456 | 21 | 1 (5%) | 0 | 0 | 0 | |
| *Act42A* | 104731 | 19 | 2 (11%) | 0 | 0 | 0 | |
| *Act5C* | 7139 | 12 | 0 | 0 | 0 | 0 | |
| *Act5C* | 101438 | 19 | 1 (5%) | 1 (5%) | 0 | 0 | |
| *Act5C* | 102129 | 12 | 0 | 0 | 0 | 0 | |
| *Act79B* | 102588 | 13 | 0 | 0 | 0 | 0 | |
| *Act87E* | 102480 | 12 | 1 (8%) | 0 | 0 | 0 | |
| *Act88F* | 9780 | 14 | 1 (7%) | 0 | 0 | 0 | |
| *Actn* | 7760 | 24 | 1 (4%) | 0 | 1 (4%) | 0 | |
| *Actn* | 7761 | 24 | 0 | 0 | 0 | 0 | |
| *Actn* | 7762 | 26 | 5 (19%) | 0 | 0 | 0 | |
| *Actn* | 34874 | 22 | 1 (5%) | 0 | 0 | 0 | |
| *Actn* | 110719 | 26 | 0 | 0 | 0 | 0 | |
| *Arp2* | 29944 | 19 | 3 (16%) | 0 | 0 | 0 | |
| *Arp2* | 101999 | 18 | 0 | 0 | 0 | 0 | |
| *Arp3* | 35260 | 20 | 1 (5%) | 0 | 0 | 0 | |
| *atk* | 64917 | 27 | 2 (7%) | 3 (11%) | 0 | 0 | |
| *Blimp* | 34978 | 23 | 6 (26%) | 3 (13%) | 0 | 0 | U |
| *Bsg* | 2789 | 28 | 0 | 0 | 0 | 0 | |
| *Bsg* | 43306 | 23 | 2 (9%) | 0 | 0 | 0 | |
| *Bsg* | 43307 | 25 | 0 | 0 | 0 | 0 | |
| *Bsg* | 52110 | 23 | 5 (22%) | 0 | 0 | 0 | |
| *Bsg* | 105293 | 23 | 2 (9%) | 0 | 0 | 0 | |
| *Cad86C* | 21327 | 3 | 0 | 0 | 0 | 0 | |
| *Cad99C* | 3739 | 7 | 0 | 2 (29%) | 0 | 0 | |
| *Cad99C* | 3739 | 5 | 0 | 1 (20%) | 0 | 0 | |
| *Cad99C* | 27212 | 2 | 0 | 1 (50%) | 0 | 0 | |
| *Cad99C* | 27212 | 4 | 0 | 1 (25%) | 0 | 0 | |
| *Cad99C* | 27212 | 8 | 0 | 1 (12%) | 0 | 0 | |

| Gene | RNAi line (BDSC or VDRC stock #) | Hemi-segments | Undulated cap | Cap–CA detachment | Cap thinning | Reduced cap/ligament ratio | Positive |
|---|---|---|---|---|---|---|---|
| *Cad99C* | 27216 | 11 | 3 (27%) | 0 | 0 | 0 | |
| *Cad99C* | 27510 | 26 | 4 (15%) | 0 | 0 | 0 | |
| *Cad99C* | 27711 | 27 | 3 (11%) | 0 | 0 | 0 | |
| *CadN2* | 47538 | 3 | 0 | 1 (33%) | 0 | 0 | |
| *cals* | 105111 | 7 | 1 (14%) | 0 | 0 | 0 | |
| *cathD* | 28978 | 21 | 0 | 1 (5%) | 0 | 0 | |
| *cathD* | 55178 | 21 | 3 (14%) | 0 | 0 | 1 (5%) | |
| *CG3168* | 29301 | 28 | 9 (32%) | 3 (11%) | 0 | 0 | U |
| *CG3168* | 48010 | 25 | 2 (8%) | 0 | 0 | 0 | |
| *CG3168* | 48011 | 25 | 0 | 1 (4%) | 0 | 0 | |
| *CG4096* | 109025 | 18 | 2 (11%) | 0 | 0 | 0 | |
| *Chb* | 26051 | 17 | 4 (24%) | 0 | 0 | 0 | |
| *Chb* | 108620 | 24 | 1 (4%) | 1 (4%) | 0 | 0 | |
| *chc* | 27530 | 30 | 0 | 0 | 0 | 0 | |
| *chc* | 34742 | 26 | 4 (15%) | 0 | 0 | 0 | |
| *Cheerio* | 107451 | 20 | 5 (25%) | 0 | 0 | 0 | U |
| *clos* | 32122 | 23 | 1 (4%) | 0 | 0 | 0 | |
| *crumbs* | 27697 | 22 | 0 | 1 (5%) | 0 | 0 | |
| *crumbs* | 34999 | 23 | 1 (4%) | 2 (9%) | 0 | 0 | |
| *crumbs* | 38373 | 20 | 5 (25%) | 1 (5%) | 0 | 0 | U |
| *crumbs* | 38903 | 21 | 1 (5%) | 0 | 0 | 0 | |
| *crumbs* | 39177 | 23 | 2 (9%) | 1 (4%) | 0 | 0 | |
| *CtsB* | 28602 | 25 | 1 (4%) | 0 | 0 | 0 | |
| *CtsB* | 33953 | 27 | 0 | 1 (4%) | 0 | 0 | |
| *Ddr* | 29720 | 2 | 0 | 0 | 0 | 0 | |
| *Ddr* | 51719 | 35 | 0 | 0 | 0 | 0 | |
| *Dg* | 10702 | 27 | 1 (4%) | 1 (4%) | 0 | 0 | |
| *Dg* | 34895 | 31 | 1 (3%) | 0 | 0 | 0 | |
| *dlt* | 41876 | 22 | 3 (14%) | 1 (5%) | 0 | 0 | |
| *dpy* | 25823 | 28 | 0 | 2 (7%) | 0 | 0 | |
| *ds* | 4313 | 22 | 0 | 0 | 0 | 0 | |
| *ds* | 36219 | 20 | 0 | 0 | 0 | 0 | |
| *eys* | 32599 | 5 | 0 | 0 | 0 | 0 | |
| *Fak56D* | 29323 | 23 | 3 (13%) | 0 | 0 | 0 | |
| *Fak56D* | 33617 | 22 | 0 | 0 | 0 | 0 | |
| *Fak56D* | 35357 | 20 | 3 (15%) | 0 | 0 | 0 | |
| *Fak56D* | 44075 | 23 | 6 (26%) | 0 | 0 | 0 | U |

| Gene | RNAi line (BDSC or VDRC stock #) | Hemi-segments | Undulated cap | Cap–CA detachment | Cap thinning | Reduced cap/ligament ratio | Positive |
|---|---|---|---|---|---|---|---|
| *Fermitin 1 §* | 46494 | 17 | 0 | 5 (29%) | 16 (94%) | — | D,T |
| *Fermitin 2 §* | 37009 | 26 | 6 (23%) | 0 | 12 (46%) | — | U,T |
| *Fermitin 2 §* | 37010 | 21 (+1) | 9 (41%) | 3 (14%) | 21 (100%) | — | U,T |
| *ft* | 9396 | 19 | 3 (16%) | 1 (5%) | 0 | 0 | |
| *ft* | 108863 | 16 | 0 | 1 (6%) | 0 | 1 (6%) | |
| *Fur1* | 25873 | 20 | 0 | 0 | 0 | 0 | |
| *Fur1* | 41914 | 23 | 0 | 3 (13%) | 0 | 0 | |
| *Fur1* | 42481 | 20 | 3 (15%) | 0 | 0 | 0 | |
| *fw* | 39575 | 27 | 1 (4%) | 0 | 0 | 0 | |
| *fw* | 106656 | 24 | 1 (4%) | 0 | 1 (4%) | 0 | |
| *Gfrl* | 103213 | 16 | 0 | 1 (6%) | 0 | 0 | |
| *Gfrl* | 103523 | 3 | 1 (33%) | 0 | 0 | 0 | |
| *glt* | 15428 | 35 | 0 | 0 | 0 | 0 | |
| *if §* | 5600 | 12 (+2) | 2 (14%) | 0 | 0 | — | |
| *if §* | 27544 | 20 (+3) | 3 (13%) | 1 (5%) | 0 | — | |
| *if §* | 38958 | 17 | 0 | 1 (6%) | 0 | — | |
| *if §* | 44885 | 14 (+13) | 13 (48%) | 0 | 0 | — | U |
| *if §* | 65346 | 16 (+5) | 5 (24%) | 1 (6%) | 0 | — | |
| *if §* | 100770 | 19 (+6) | 6 (24%) | 0 | 3 (16%) | — | U,T |
| *kirre* | 109585 | 2 | 0 | 0 | 0 | 0 | |
| *kug* | 110390 | 12 | 1 (8%) | 0 | 0 | 0 | |
| *l(2)gl* | 31089 | 12 | 2 (17%) | 1 (8%) | 0 | 0 | |
| *l(2)gl* | 38206 | 24 | 7 (29%) | 1 (4%) | 0 | 0 | U |
| *l(2)gl* | 38989 | 24 | 4 (17%) | 1 (4%) | 0 | 0 | |
| *LanB2* | 104013 | 34 | 0 | 0 | 0 | 0 | |
| *Lasp* | 26305 | 26 | 0 | 0 | 0 | 0 | |
| *Lcp1* | 106030 | 10 | 0 | 1 (10%) | 0 | 0 | |
| *Lcp2* | 12537 | 8 | 0 | 0 | 0 | 0 | |
| *Lcp3* | 12540 | 22 | 1 (5%) | 2 (9%) | 0 | 0 | |
| *Liprin-a* | 53868 | 23 | 1 (4%) | 0 | 0 | 0 | |
| *Loh* | 31020 | 18 | 1 (6%) | 0 | 0 | — | |
| *Loxl1* | 17259, 16619 | 26 | 6 (23%) | 0 | 0 | 0 | U |
| *mew §* | 44890 | 54 (+1) | 2 (4%) | 0 | 0 | — | |
| *mew §* | 109608 | 42 | 1 (2%) | 3 (7%) | 0 | — | |
| *mlt* | 105480 | 26 | 4 (15%) | 0 | 0 | 0 | |
| *MMP1* | 31489 | 32 | 0 | 1 (3%) | 0 | 0 | |
| *MMP1* | 101505 | 27 | 0 | 0 | 0 | 0 | |

| Gene | RNAi line (BDSC or VDRC stock #) | Hemi-segments | Undulated cap | Cap–CA detachment | Cap thinning | Reduced cap/ligament ratio | Positive |
|---|---|---|---|---|---|---|---|
| *MMP1 + MMP2* | 10505, 65935 | 22 | 1 (5%) | 0 | 0 | 0 | |
| *MMP1 + MMP2* | 107883, 31489 | 23 | 1 (4%) | 0 | 0 | 0 | |
| *MMP2* | 31371 | 26 | 2 (8%) | 1 (4%) | 0 | 0 | |
| *MMP2* | 61309 | 29 | 3 (10%) | 1 (3%) | 0 | 0 | |
| *MMP2* | 65935 | 23 | 3 (13%) | 0 | 0 | 0 | |
| *MMP2* | 107888 | 27 | 0 | 1 (4%) | 0 | 0 | |
| *Moe* | 31135 | 26 | 6 (23%) | 0 | 0 | 0 | U |
| *Moe* | 31872 | 22 | 1 (5%) | 0 | 0 | 0 | |
| *Moe* | 33936 | 19 | 4 (21%) | 0 | 0 | 0 | |
| *mspo* | 107608 | 4 | 0 | 0 | 0 | 0 | |
| *Muc12Ea* | 50435 | 7 | 2 (29%) | 2 (29%) | 0 | 0 | |
| *Muc12Ea* | 50435 | 6 | 0 | 0 | 0 | 0 | |
| *Muc55B* | 104747 | 5 | 0 | 1 (20%) | 0 | 0 | |
| *mulet* | 104780 | 28 | 6 (21%) | 0 | 0 | 0 | |
| *Mur89F* | 34794 | 12 | 0 | 0 | 0 | 0 | |
| *Mur89F* | 100128 | 2 | 0 | 0 | 0 | 0 | |
| *Ndg* | 13280 | 3 | 0 | 0 | 0 | 0 | |
| *nolo (CG31619)* | 33102 | 2 | 0 | 0 | 0 | 0 | |
| *nolo (CG31619)* | 104736 | 2 | 0 | 0 | 0 | 0 | |
| *obst-A* | 12935 | 6 | 0 | 1 (17%) | 0 | 0 | |
| *obst-A* | 102591 | 2 | 0 | 0 | 0 | 0 | |
| *Pak* | 35141 | 19 | 0 | 0 | 0 | 0 | |
| *Pak* | 41714 | 24 | 8 (33%) | 0 | 0 | 0 | U |
| *Parvin §* | 11670 | 16 | 0 | 0 | 0 | — | |
| *Parvin §* | 105356 | 16 | 0 | 0 | 0 | — | |
| *Patj* | 26282 | 24 | 7 (29%) | 0 | 0 | 0 | U |
| *Patj* | 35747 | 28 | 5 (18%) | 1 (4%) | 0 | 0 | |
| *Patj* | 38193 | 26 | 8 (31%) | 1 (4%) | 0 | 0 | U |
| *Pax* | 25853 | 18 | 1 (6%) | 0 | 0 | 2 (11%) | |
| *Pax* | 107789 | 20 | 1 (5%) | 0 | 0 | 0 | |
| *Plod* | 45484 | 56 | 0 | 0 | 0 | 0 | |
| *Plod* | 45484 | 11 | 0 | 1 (9%) | 0 | 0 | |
| *pyd* | 38863 | 3 | 0 | 0 | 0 | 0 | |
| *Rap1* | 33437 | 24 | 0 | 0 | 0 | 2 (8%) | |
| *rhea* | 28950 | 16 | — | 7 (44%) | 10 (62%) | — | D,T |
| *rhea* | 32999 | 14 | 0 | 0 | 11 (79%) | 11 (79%) | T |
| *rhea §* | 40399 | 21 (+2) | 2 (9%) | 1 (5%) | 7 (33%) | — | T |

| Gene | RNAi line (BDSC or VDRC stock #) | Hemi-segments | Undulated cap | Cap–CA detachment | Cap thinning | Reduced cap/ligament ratio | Positive |
|---|---|---|---|---|---|---|---|
| *rhea §* | 40400 | 17 (+1) | 2 (11%) | 0 | 3 (18%) | — | T |
| *sdc* | 13322 | 23 | 2 (9%) | 1 (4%) | 0 | 0 | |
| *sdc* | 51723 | 26 | 2 (8%) | 1 (4%) | 0 | 0 | |
| *sdc* | 107320 | 27 | 3 (11%) | 5 (19%) | 1 (4%) | 0 | |
| *Sec23* | 24552 | 30 | 0 | 0 | 0 | 0 | |
| *Sec23* | 32365 | 22 | 6 (27%) | 1 (5%) | 0 | 0 | U |
| *Sec23* | 110568 | 29 | 0 | 2 (7%) | 2 (7%) | 0 | |
| *Sec8* | 45032 | 23 | 0 | 0 | 0 | 0 | |
| *Sec8* | 105653 | 26 | 0 | 0 | 0 | 0 | |
| *sgs1* | 21207 | 13 | 0 | 1 (8%) | 0 | 0 | |
| *stck §* | 52537 | 16 | 0 | 0 | 0 | — | |
| *stck §* | 52538 | 20 | 0 | 0 | 1 (5%) | — | |
| *stck §* | 100582 | 22 | 2 (9%) | 0 | 0 | — | |
| *Tensin* | 22823 | 23 | 0 | 0 | 0 | 0 | |
| *Tensin* | 330366 | 23 | 1 (4%) | 2 (9%) | 0 | 0 | |
| *Timp* | 15373 | 22 | 0 | 0 | 0 | 0 | |
| *Timp* | 109427 | 23 | 5 (22%) | 0 | 0 | 0 | |
| *Transglutaminase (Tg)* | 103601 | 4 | 0 | 0 | 0 | 0 | |
| *tsp* | 7535 | 19 | 0 | 7 (37%) | 0 | 0 | D |
| *tsp* | 100721 | 17 | 0 | 8 (47%) | 0 | 0 | D |
| *TwdlE* | 107483 | 56 | 0 | 1 (2%) | 0 | 0 | |
| *TwdlF* | 18903 | 12 | 1 (8%) | 0 | 0 | 0 | |
| *TwdlG* | 100335 | 1 | 0 | 0 | 0 | 0 | |
| *TwdlJ* | 103260 | 11 | 0 | 1 (9%) | 0 | 0 | |
| *TwdlT* | 41335 | 8 | 0 | 0 | 0 | 0 | |
| *TwdlW* | 101966 | 7 | 0 | 1 (14%) | 0 | 0 | |
| *Vinc* | 25965 | 24 | 3 (12%) | 0 | 0 | 0 | |
| *Vinc* | 34586 | 25 | 2 (8%) | 0 | 0 | 0 | |
| *Vinc* | 41959 | 31 | 9 (29%) | 0 | 0 | 0 | U |
| *Vinc §* | 34584 | 22 | 0 | 0 | 0 | — | |
| *Vinc §* | 34586 | 17 (+3) | 3 (15%) | 0 | 0 | — | |
| *Vinc §* | 105956 | 18 (+3) | 3 (14%) | 0 | 0 | — | |
| *Yki* | 111001 | 24 | 3 (12%) | 0 | 0 | 0 | |
| *Zasp52* | 31561 | 21 | 1 (5%) | 1 (5%) | 0 | 0 | |
| *Zasp52* | 58198 | 23 | 3 (13%) | 0 | 0 | 0 | |
| *Zyxin* | 21610 | 24 | 1 (4%) | 0 | 0 | 0 | |
| *Ddr* | 51719 | 49 | 0 | 1 (2%) | 0 | 0 | |

| Gene | RNAi line<br>(BDSC or VDRC stock #) | Hemi-<br>segments | Undulated cap | Cap–CA detachment | Cap thinning | Reduced cap/ligament ratio | Positive |
|---|---|---|---|---|---|---|---|
| *LanB2* | 104013 | 42 | 0 | 1 (2%) | 0 | 0 | |
| *Plod* | 45484 | 49 | 0 | 0 | 0 | 0 | |
| *TwdlE* | 107483 | 28 | 0 | 0 | 0 | 0 | |
| *TwdlW* | 101966 | 7 | 0 | 0 | 0 | 0 | |
| *dpy* | 25823 | 28 | 0 | 2 (7%) | 0 | 0 | |
| *glt* | 15428 | 35 | 0 | 0 | 0 | 0 | |

Columns and the Positive criterion as in Table 1, with comparison to the *en* > Luc RNAi control.

§ (after the gene name) Genotypes analyzed in Table 4: detachment and thinning were counted in measured hemisegments; undulation also includes additional hemisegments shown in parentheses in the Hemisegments column.

Genes with at least two positive lines for the same phenotype: undulation, *Patj*, *if* and *Fermitin 2*; cap thinning, *rhea* and *Fermitin 2*; cap–CA detachment, *tsp*.

**Totals (excluding the control and *prc*): 170 RNAi lines or line combinations, 90 genes, 176 experiments and 3,463 hemisegments.**

### Table 4. Cap and ligament dimensions after LCh5-directed (*en*-Gal4) knockdown of integrin pathway components

| Gene | RNAi line (BDSC or VDRC stock #) | Larvae | Hemi-segments | Organ length (µm) | Cap length (µm) | L+SenU length (µm) | C/(L+SenU) | Below 1.1 | Reduced ratio (size-matched) | Elongated ligament (size-matched) | L+SenU, fold of size-matched control | Positive |
|---|---|---|---|---|---|---|---|---|---|---|---|---|
| *Luc (control)* | 31603 | 13 | 45 | 127.7 ± 21.0 | 79.4 ± 17.3 | 48.3 ± 5.7 | 1.64 ± 0.30 | 0/45 (0%) | 1/45 (2%) | 1/45 (2%) | 1.00 | |
| *if* | 100770 | 8 | 19 | 94.6 ± 11.6 | 55.0 ± 7.1 | 39.6 ± 5.6 | 1.40 ± 0.15 | 0/19 (0%) | 0/19 (0%) | 0/19 (0%) | 0.95 | |
| *if* | 27544 | 7 | 20 | 81.2 ± 6.8 | 36.5 ± 4.9 | 44.7 ± 5.8 | 0.83 ± 0.17 | 20/20 (100%) | 13/20 (65%) | 6/20 (30%) | 1.14 | R,E |
| *if* | 38958 | 9 | 17 | 83.1 ± 17.4 | 46.2 ± 13.2 | 36.9 ± 6.5 | 1.26 ± 0.29 | 4/17 (24%) | 1/17 (6%) | 1/17 (6%) | 0.93 | |
| *if* | 44885 | 7 | 14 | 72.9 ± 11.6 | 31.1 ± 9.8 | 41.7 ± 3.6 | 0.75 ± 0.20 | 13/14 (93%) | 10/14 (71%) | 1/14 (7%) | 1.11 | R |
| *if* | 5600 | 4 | 12 | 81.9 ± 16.4 | 45.5 ± 12.4 | 36.4 ± 4.8 | 1.24 ± 0.24 | 5/12 (42%) | 0/12 (0%) | 0/12 (0%) | 0.93 | |
| *if* | 65346 | 7 | 16 | 85.1 ± 13.1 | 47.6 ± 10.6 | 37.5 ± 4.4 | 1.27 ± 0.26 | 5/16 (31%) | 1/16 (6%) | 0/16 (0%) | 0.94 | |
| *rhea* | 28950 | 5 | 14 | 67.7 ± 5.5 | 33.5 ± 4.3 | 34.2 ± 3.2 | 0.99 ± 0.16 | 10/14 (71%) | 1/14 (7%) | 0/14 (0%) | 0.94 | |
| *rhea* | 40399 | 7 | 21 | 79.6 ± 16.3 | 37.4 ± 14.8 | 42.2 ± 4.2 | 0.88 ± 0.34 | 17/21 (81%) | 16/21 (76%) | 2/21 (10%) | 1.09 | R |
| *rhea* | 40400 | 7 | 17 | 65.4 ± 4.7 | 30.2 ± 4.8 | 35.3 ± 4.5 | 0.88 ± 0.23 | 14/17 (82%) | 6/17 (35%) | 0/17 (0%) | 0.98 | R |
| *stck* | 100582 | 8 | 22 | 126.9 ± 15.0 | 59.0 ± 10.9 | 67.9 ± 18.4 | 0.95 ± 0.38 | 14/22 (64%) | 16/22 (73%) | 16/22 (73%) | 1.41 | R,E |
| *stck* | 52537 | 7 | 16 | 147.4 ± 29.7 | 53.3 ± 10.8 | 94.0 ± 21.3 | 0.58 ± 0.09 | 16/16 (100%) | 16/16 (100%) | 16/16 (100%) | 1.80 | R,E |
| *stck* | 52538 | 8 | 20 | 135.3 ± 37.4 | 45.9 ± 9.0 | 89.5 ± 30.2 | 0.54 ± 0.11 | 20/20 (100%) | 20/20 (100%) | 20/20 (100%) | 1.80 | R,E |
| *Parvin* | 105356 | 6 | 16 | 115.3 ± 20.9 | 40.3 ± 8.0 | 75.0 ± 17.0 | 0.56 ± 0.16 | 16/16 (100%) | 16/16 (100%) | 16/16 (100%) | 1.63 | R,E |
| *Parvin* | 11670 | 7 | 16 | 139.2 ± 26.8 | 46.8 ± 4.3 | 92.4 ± 25.1 | 0.54 ± 0.14 | 16/16 (100%) | 16/16 (100%) | 16/16 (100%) | 1.83 | R,E |
| *Fermitin 1* | 46494 | 7 | 17 | 122.0 ± 24.6 | 81.6 ± 20.9 | 40.5 ± 5.6 | 2.02 ± 0.44 | 0/17 (0%) | 0/17 (0%) | 0/17 (0%) | 0.86 | |
| *Fermitin 2* | 37009 | 8 | 26 | 90.9 ± 14.0 | 53.5 ± 10.1 | 37.4 ± 6.0 | 1.45 ± 0.25 | 4/26 (15%) | 2/26 (8%) | 0/26 (0%) | 0.91 | |
| *Fermitin 2* | 37010 | 8 | 21 | 100.3 ± 6.6 | 65.6 ± 5.8 | 34.7 ± 2.6 | 1.90 ± 0.21 | 0/21 (0%) | 0/21 (0%) | 0/21 (0%) | 0.81 | |
| *Vinc* | 105956 | 7 | 18 | 109.7 ± 22.8 | 66.0 ± 16.0 | 43.7 ± 8.2 | 1.51 ± 0.25 | 1/18 (6%) | 0/18 (0%) | 0/18 (0%) | 0.98 | |
| *Vinc* | 25965 | 9 | 22 | 95.3 ± 11.8 | 56.4 ± 8.6 | 38.9 ± 6.1 | 1.48 ± 0.32 | 1/22 (5%) | 0/22 (0%) | 0/22 (0%) | 0.93 | |
| *Vinc* | 34584 | 9 | 22 | 113.7 ± 24.4 | 64.0 ± 17.7 | 49.7 ± 7.8 | 1.28 ± 0.23 | 6/22 (27%) | 5/22 (23%) | 4/22 (18%) | 1.09 | R,E |
| *Vinc* | 34586 | 7 | 17 | 116.6 ± 27.0 | 65.1 ± 15.9 | 51.5 ± 11.6 | 1.26 ± 0.12 | 1/17 (6%) | 7/17 (41%) | 6/17 (35%) | 1.12 | R,E |
| *Vinc* | 41959 | 9 | 21 | 93.9 ± 15.8 | 53.1 ± 12.3 | 40.8 ± 4.7 | 1.30 ± 0.22 | 2/21 (10%) | 0/21 (0%) | 0/21 (0%) | 0.98 | |
| *mew* | 109608 | 8 | 42 | 108.3 ± 10.6 | 66.6 ± 8.2 | 41.7 ± 4.0 | 1.60 ± 0.19 | 0/42 (0%) | 0/42 (0%) | 0/42 (0%) | 0.94 | |
| *mew* | 44890 | 10 | 54 | 103.6 ± 15.0 | 61.1 ± 11.8 | 42.5 ± 4.6 | 1.44 ± 0.20 | 4/54 (7%) | 1/54 (2%) | 1/54 (2%) | 0.98 | |

Columns as in Table 2, with size-specific limits derived from the *en* > Luc RNAi control.

### Table 5. Cap and ligament dimensions after ligament-cell-specific (*repo*-Gal4) knockdown

| Gene | RNAi line (BDSC or VDRC stock #) | Larvae | Hemi-segments | Organ length (µm) | Cap length (µm) | L+SenU length (µm) | C/(L+SenU) | Below 1.1 | Reduced ratio (size-matched) | Elongated ligament (size-matched) | L+SenU, fold of size-matched control | Positive |
|---|---|---|---|---|---|---|---|---|---|---|---|---|
| ***Reference: en-Gal4 > Luc RNAi (Table 3)*** | | | | | | | | | | | | |
| *Luc (en-Gal4 control)* | 31603 | 13 | 45 | 127.7 ± 21.0 | 79.4 ± 17.3 | 48.3 ± 5.7 | 1.64 ± 0.30 | 0/45 (0%) | 1/45 (2%) | 1/45 (2%) | 1.00 | |
| | | | | | | ***repo-Gal4*** | | | | | | |
| *mys* | 29619 | 6 | 15 | 129.1 ± 25.6 | 38.3 ± 5.6 | 90.8 ± 25.4 | 0.46 ± 0.16 | 15/15 (100%) | 15/15 (100%) | 15/15 (100%) | 1.87 | R,E |
| *mys* | 33642 | 6 | 13 | 152.2 ± 23.9 | 37.7 ± 2.9 | 114.5 ± 21.8 | 0.34 ± 0.06 | 13/13 (100%) | 13/13 (100%) | 13/13 (100%) | 2.16 | R,E |
| *if* | 27544 | 5 | 15 | 112.0 ± 9.1 | 50.7 ± 8.5 | 61.2 ± 7.9 | 0.85 ± 0.21 | 13/15 (87%) | 14/15 (93%) | 13/15 (87%) | 1.35 | R,E |
| *if* | 44885 | 7 | 19 | 103.3 ± 9.6 | 42.7 ± 6.5 | 60.6 ± 6.1 | 0.71 ± 0.12 | 19/19 (100%) | 19/19 (100%) | 19/19 (100%) | 1.39 | R,E |
| *rhea* | 32999 | 5 | 13 | 118.2 ± 27.9 | 37.0 ± 9.1 | 81.1 ± 21.3 | 0.47 ± 0.09 | 13/13 (100%) | 13/13 (100%) | 13/13 (100%) | 1.75 | R,E |
| *rhea* | 40399 | 7 | 15 | 153.9 ± 29.9 | 47.2 ± 15.6 | 106.7 ± 25.3 | 0.46 ± 0.18 | 15/15 (100%) | 15/15 (100%) | 15/15 (100%) | 2.00 | R,E |
| *stck* | 52538 | 6 | 17 | 118.5 ± 25.7 | 43.3 ± 6.9 | 75.2 ± 19.9 | 0.60 ± 0.12 | 17/17 (100%) | 17/17 (100%) | 16/17 (94%) | 1.62 | R,E |
| *Fermitin 2* | 37010 | 7 | 18 | 118.1 ± 24.5 | 65.9 ± 13.1 | 52.2 ± 12.8 | 1.29 ± 0.19 | 3/18 (17%) | 6/18 (33%) | 7/18 (39%) | 1.12 | R,E |

Columns as in Table 2.

### Table 6. Cap and ligament dimensions after cap-cell-specific (*β3tub-cap*-Gal4) knockdown, with the paired *repo*-Gal4 value

| Gene | RNAi line (BDSC or VDRC stock #) | Larvae | Hemi-segments | Organ length (µm) | Cap length (µm) | L+SenU length (µm) | C/(L+SenU) | Below 1.1 | Reduced ratio (size-matched) | Elongated ligament (size-matched) | L+SenU, fold of size-matched control | Defects noted | Same line under repo-Gal4: C/(L+SenU); p |
|---|---|---|---|---|---|---|---|---|---|---|---|---|---|
| ***Reference: en-Gal4 > Luc RNAi (Table 3)*** | | | | | | | | | | | | | |
| *Luc (en-Gal4 control)* | 31603 | 13 | 45 | 127.7 ± 21.0 | 79.4 ± 17.3 | 48.3 ± 5.7 | 1.64 ± 0.30 | 0/45 (0%) | 1/45 (2%) | 1/45 (2%) | 1.00 | — | — |
| ***β3tub-cap-Gal4*** | | | | | | | | | | | | | |
| *mys* | 29619 | 6 | 15 | 134.0 ± 46.1 | 89.3 ± 40.3 | 44.7 ± 6.1 | 1.93 ± 0.63 | 0/15 (0%) | 0/15 (0%) | 0/15 (0%) | 0.90 | 2 × undulated cap | 0.46; p = 3e-6 |
| *mys* | 33642 | 6 | 16 | 104.2 ± 15.6 | 66.6 ± 12.7 | 37.5 ± 4.8 | 1.78 ± 0.28 | 0/16 (0%) | 0/16 (0%) | 0/16 (0%) | 0.86 | — | 0.34; p = 6e-6 |
| *if* | 44885 | 6 | 16 | 118.9 ± 39.0 | 74.7 ± 32.9 | 44.2 ± 7.3 | 1.65 ± 0.48 | 0/16 (0%) | 0/16 (0%) | 0/16 (0%) | 0.95 | — | 0.71; p = 5e-7 |
| *rhea* | 32999 | 4 | 12 | 98.0 ± 26.8 | 60.6 ± 21.1 | 37.4 ± 7.8 | 1.62 ± 0.42 | 0/12 (0%) | 0/12 (0%) | 0/12 (0%) | 0.88 | 1 × detached cap | 0.47; p = 2e-5 |
| *rhea* | 40399 | 6 | 14 | 125.4 ± 12.1 | 79.8 ± 11.4 | 45.6 ± 3.5 | 1.76 ± 0.28 | 0/14 (0%) | 0/14 (0%) | 0/14 (0%) | 0.95 | — | 0.46; p = 5e-6 |
| *stck* | 52538 | 7 | 19 | 116.6 ± 37.2 | 76.2 ± 31.4 | 40.4 ± 6.6 | 1.84 ± 0.48 | 0/19 (0%) | 0/19 (0%) | 0/19 (0%) | 0.88 | — | 0.60; p = 3e-7 |
| *Fermitin 1* | 46494 | 6 | 17 | 94.4 ± 19.8 | 55.6 ± 16.1 | 38.9 ± 4.5 | 1.42 ± 0.25 | 0/17 (0%) | 0/17 (0%) | 0/17 (0%) | 0.93 | — | not tested |
| *Fermitin 2* | 37010 | 9 | 25 | 99.7 ± 20.8 | 61.6 ± 17.2 | 38.1 ± 4.4 | 1.60 ± 0.28 | 0/25 (0%) | 0/25 (0%) | 0/25 (0%) | 0.89 | 3 × undulated cap | 1.29; p = 1e-4 |

Columns as in Table 2. Defects noted: hemisegments with undulated or detached cap cells recorded on the measurement sheet. The final column gives the mean C/(L+SenU) obtained with the same RNAi line under *repo*-Gal4 (Table 4) and the two-sided Mann–Whitney p value for the comparison between drivers.

### Table 7. Conditional (tub-Gal80ts) knockdown of *mys* in ligament cells across temperature regimes

| Genotype and temperature regime | n | L+SenU (µm) | Cap (µm) | Total GFP length (µm) | Ligament fraction | C/(L+SenU) equivalent | p vs control |
|---|---|---|---|---|---|---|---|
| *repo*-Gal4 > tub-Gal80ts, GFP — 18°C | 22 | 65.4 ± 5.3 | 348.7 ± 26.0 | 414.1 ± 28.1 | 0.158 ± 0.012 | 5.32 | reference |
| *repo*-Gal4 > tub-Gal80ts, GFP; *mys*-RNAi — 18°C | 24 | 76.7 ± 9.8 | 334.1 ± 42.6 | 410.9 ± 46.4 | 0.188 ± 0.022 | 4.32 | 2e-06 |
| *repo*-Gal4 > *mys*-RNAi (no Gal80ts) — 18°C | 24 | 376.2 ± 37.8 | 67.0 ± 9.9 | 443.2 ± 40.7 | 0.848 ± 0.020 | 0.18 | 7e-09 |
| *repo*-Gal4 > tub-Gal80ts, GFP — 29°C | 24 | 59.5 ± 4.6 | 329.6 ± 26.1 | 389.1 ± 29.1 | 0.153 ± 0.009 | 5.53 | reference |
| *repo*-Gal4 > tub-Gal80ts, GFP; *mys*-RNAi — 29°C | 25 | 390.7 ± 22.2 | 68.7 ± 14.5 | 459.4 ± 29.1 | 0.851 ± 0.026 | 0.17 | 2e-09 |
| *repo*-Gal4 > tub-Gal80ts, GFP — 18°C → 29°C | 24 | 61.6 ± 4.5 | 311.7 ± 25.2 | 373.3 ± 26.1 | 0.166 ± 0.014 | 5.04 | reference |
| *repo*-Gal4 > tub-Gal80ts, GFP; *mys*-RNAi — 18°C → 29°C | 24 | 140.1 ± 33.3 | 237.2 ± 44.9 | 377.4 ± 36.4 | 0.373 ± 0.089 | 1.68 | 3e-09 |
| *repo*-Gal4 > tub-Gal80ts, GFP; *mys*-RNAi — 29°C → 18°C | 24 | 397.6 ± 47.4 | 74.3 ± 21.2 | 471.9 ± 50.7 | 0.843 ± 0.043 | 0.19 | 3e-09 |
| *repo*-Gal4 > tub-Gal80ts, GFP; *mys*-RNAi — 18→29°C, shifted 14 h AEL | 21 | 295.7 ± 49.3 | 123.4 ± 29.1 | 419.1 ± 37.2 | 0.703 ± 0.076 | 0.42 | 1e-08 |
| *repo*-Gal4 > tub-Gal80ts, GFP; *mys*-RNAi — 18→29°C, shifted 12 h AEL | 19 | 343.4 ± 54.8 | 92.3 ± 41.0 | 435.8 ± 26.0 | 0.786 ± 0.098 | 0.27 | 3e-08 |
| *repo*-Gal4 / Canton S — 29°C, first instar | 20 | 29.3 ± 4.0 | 39.1 ± 4.4 | 68.4 ± 5.9 | 0.428 ± 0.043 | 1.34 | reference |
| *repo*-Gal4 > *mys*-RNAi — 29°C, first instar | 20 | 42.3 ± 5.8 | 27.4 ± 3.0 | 69.8 ± 6.2 | 0.605 ± 0.043 | 0.65 | 7e-08 |

*mys RNAi* #29619 was used in the experiments. Mean ± SD; n = organs measured. Rows 1–10 are third-instar larvae; the final two rows are first instar. Ligament fraction = L+SenU length as a fraction of total labelled organ length. C/(L+SenU) equivalent = (1 − fraction)/fraction, placing these regimes on the same scale as Tables 3–5. p vs control = two-sided Mann–Whitney test against the temperature-matched driver control marked “reference”; the 29°C → 18°C and staged shift regimes are tested against the 18°C → 29°C control.
